# TET1 Catalytic Activity is Required for Reprogramming of Imprinting Control Regions and Patterning of Sperm-Specific Hypomethylated Regions

**DOI:** 10.1101/2023.02.21.529426

**Authors:** Rexxi D. Prasasya, Blake A. Caldwell, Zhengfeng Liu, Songze Wu, Nicolae A. Leu, Johanna M. Fowler, Steven A. Cincotta, Diana J. Laird, Rahul M. Kohli, Marisa S. Bartolomei

**Author notes:** Co-lead contacts (Rahul M. Kohli, Marisa S. Bartolomei). Correspondence (MSB) and (RMK).

## Abstract

DNA methylation erasure is required for mammalian primordial germ cell reprogramming. TET enzymes iteratively oxidize 5-methylcytosine to generate 5-hyroxymethylcytosine (5hmC), 5-formylcytosine, and 5-carboxycytosine to facilitate active genome demethylation. Whether these bases are required to promote replication-coupled dilution or activate base excision repair during germline reprogramming remains unresolved due to the lack of genetic models that decouple TET activities. Here, we generated two mouse lines expressing catalytically inactive TET1 (*Tet1-HxD*) and TET1 that stalls oxidation at 5hmC (*Tet1-V*). *Tet1^-/-^*, *Tet1^V/V^*, and *Tet1^HxD/HxD^* sperm methylomes show that TET1^V^ and TET1^HxD^ rescue most *Tet1^-/-^* hypermethylated regions, demonstrating the importance of TET1’s extra-catalytic functions. Imprinted regions, in contrast, require iterative oxidation. We further reveal a broader class of hypermethylated regions in sperm of *Tet1* mutant mice that are excluded from *de novo* methylation during male germline development and depend on TET oxidation for reprogramming. Our study underscores the link between TET1-mediated demethylation during reprogramming and sperm methylome patterning.

## Introduction

DNA methylation is a major conveyer of epigenetic information in the eukaryotic genome. 5-methylcytosine (5mC) bases are found primarily within CpG dinucleotides, with 70-80% of CpGs in the mammalian genomes showing mitotically stable methylation. 5mC enrichment within enhancers and gene promoters is associated with transcriptional repression and shapes cell-specific gene expression profiles^1^. DNA methylation is also essential for maintaining genomic stability by repressing repetitive elements^2^. In the germline, DNA methylation is used to mark imprinted genes, where *cis-*regulatory elements known as imprinting control regions (ICRs) are methylated in a parent-of-origin specific manner to confer monoallelic expression of developmentally important genes^3^.

During mammalian development, there are two periods where DNA methylation is reprogrammed genome wide. The first occurs during post-fertilization embryonic development to achieve totipotency and the subsequent establishment of tissue-specific methylation patterns^1^. The second occurs in primordial germ cells (PGCs), germ cell precursors, which are specified in mammals from pluripotent somatic cells within the proximal epiblast^4, 5^. In both instances, DNA methylation erasure is achieved through a combination of two distinct mechanisms: global replication-coupled passive dilution through the suppression of maintenance DNA methyltransferase (DNMT) activity and active demethylation that is facilitated by the family of ten-eleven translocation (TET) methylcytosine dioxygenases.

TET enzymes iteratively oxidize 5mC to 5-hydroxymethylcytosine (5hmC), 5-formylcytosine (5fC), and 5-carboxylcytosine (5caC)^6, 7^. 5hmC is poorly recognized by maintenance DNMT1 complex, thus further promoting passive dilution. Alternatively, 5fC and 5caC can be excised by thymine DNA glycosylase (TDG), generating an abasic site that activates the base excision repair machinery to recover unmodified cytosine^8^. The three TET isoforms are major regulators of mammalian development. TET1 is expressed in PGCs, where it plays a role in the complete reprogramming of ICRs^9–11^ and timely activation of meiosis-associated promoters^12, 13^. TET2 and TET3, by contrast, have tissue-specific roles in somatic development^12^. Dysregulation of TET2 activity has been implicated as a major driver of hematologic malignancies^14, 15^. In zygotes, maternally deposited TET3 is responsible for active demethylation during post-fertilization epigenetic reprogramming, with the most pronounced activity in paternal pronuclei^16–18^. Inactivation of all three *Tet* genes causes early embryonic lethality due to ectopic regulation of Lefty-Nodal signaling and failed gastrulation^19^ . Finally, we and others have demonstrated the importance of active demethylation by TET proteins at gene enhancers for successful somatic cell reprogramming to pluripotency^20–22^.

The iterative modes of 5mC oxidation by TET enzymes prompt questions of whether oxidized 5mC bases have biological significance. Identifying functions for 5fC and 5caC has been challenging due to their low abundance and rapid removal by TDG^7^. Our recent work using a *Tet2* mutant with reduced efficiency for oxidizing beyond 5hmC demonstrated that generation of these higher order oxidized bases is required for a significant portion of DNA demethylation at enhancers during induced pluripotent stem cell (iPSC) formation^22^. *Tet1^-/-^* PGCs show incomplete demethylation of ICRs and meiosis- and gametogenesis-associated gene promoters^10–13^. It has long been proposed, however, that the role of TET1 in germline reprogramming is restricted to the formation of 5hmC to promote replication coupled passive dilution^23–25^. This is largely based on the rapid rate of cellular division in PGCs and the scarcity of detectable 5fC and 5caC^13, 24–26^. It is therefore unknown whether demethylation through 5fC and 5caC contributes significantly to germline reprogramming.

In addition to their role in DNA demethylation, recent studies have demonstrated the non-catalytic involvement of TET proteins in genome regulation, particularly by interacting with diverse form of epigenetic regulators. TET1 interacts with O-linked N-acetylglucosamine (OglcNAc) transferase (OGT)^27^, which in turn can regulate activities of the COMPASS family of H3K4 methylases^28^. In mouse embryonic stem cells (ESC), TET1 is recruited by Polycomb repressive complex 2 (PRC2) to bivalent promoters enriched for H3K27me3^29, 30^. More recently, Paraspeckel component 1 (PSPC1) and its cognate long noncoding RNA (lncRNA) *Neat1* have been shown to interact with TET1, together regulating the targeting of PRC2 to chromatin to maintain bivalency in the ESC genome^31^. While knockout of TET1 protein leads to methylation and chromatin changes in ESCs, expression of catalytically inactive TET1 is able to preserve H3K27me3 and H3K4me3 enrichment at bivalent promoters, further demonstrating non-catalytic importance of TET proteins^32^.

Functional studies of TET proteins have largely relied on conventional knockout mouse models, which, while highly informative, fail to distinguish between catalytic and non-catalytic TET activities. To distinguish catalytic and non-catalytic activities of TET1 and study the requirement for 5fC and 5caC generation *in vivo*, we developed two new *Tet1* genetic mouse models. The *Tet1^T^*^1642^*^V^* (*Tet1^v^*) mutant preserves 5hmC generation but has diminished oxidative activity to 5fC and 5caC, and the *Tet1^H1654Y,D1656A^*(*TET1^HxD^*) mutant expresses catalytically inactive TET1^22, 32–34^. Our results show that TET1-mediated oxidation through 5fC and 5caC is required for complete ICR reprogramming in the male germline. Additionally, methylation defects in *Tet1* mutant sperm are more extensive than previously demonstrated. Newly identified hypermethylated regions in *Tet1* mutant sperm overlap with regions that are excluded from *de novo* methylation during spermatogenesis through enrichment of H3K4me3 in prospermatogonia^35^. This finding suggests that hypomethylated regions, which are sparse within the largely hypermethylated sperm genome^36^, may originate at loci that require an active pathway for methylation erasure during germline reprogramming. Moreover, genome-wide methylation analysis of mutant sperm reveals that full length TET1^V^ and TET1^HxD^ can partially rescue hypermethylation defects observed in *Tet1^-/-^* sperm, supporting role for non-catalytic TET1 activity in germline development. The use of *Tet1* catalytic mutants to study germline epigenetic reprogramming reveals an added complexity to PGC genome regulation, in which the locus-specific modality of methylation erasure (active vs. replication coupled) appears to contribute to patterning of the sperm methylome.

## Results

### Characterization of 5hmC stalling Tet1^V^ and catalytically inactive Tet1^HxD^ mouse lines

We previously showed that a T1642V substitution in the catalytic domain of mouse TET1 (5hmC-dominant *Tet1^V^*) results in 5hmC generation without detectable 5caC in transfected HEK293T cells^37^. Similarly, simultaneous H1654Y and D1656A substitutions in the TET1 catalytic domain (catalytically inactive *Tet1^HxD^*) ablate the catalytic activity of TET1^32, 33^. To test the catalytic requirements for TET1 during mammalian germline epigenetic reprogramming, we developed two new mouse lines harboring these mutations (Figure 1A). Mice were generated through microinjections of CRISPR/Cas9-based editing reagents into one-cell mouse embryos, and founders were backcrossed for at least 4 generations (F4) onto the C57BL/6J background. Correct point mutations of the *Tet1*^V^ and *Tet1^HxD^* alleles were confirmed by Sanger sequencing and restriction fragment length polymorphism (RFLP) analysis (Figure 1B, Supplemental Figure 1A-B). Southern blotting was performed to ensure that the CRISPR/Cas9 mutagenesis did not cause chromosomal rearrangements in the *Tet1* locus (Supplemental Figure 1C-E).

**Figure 1.**
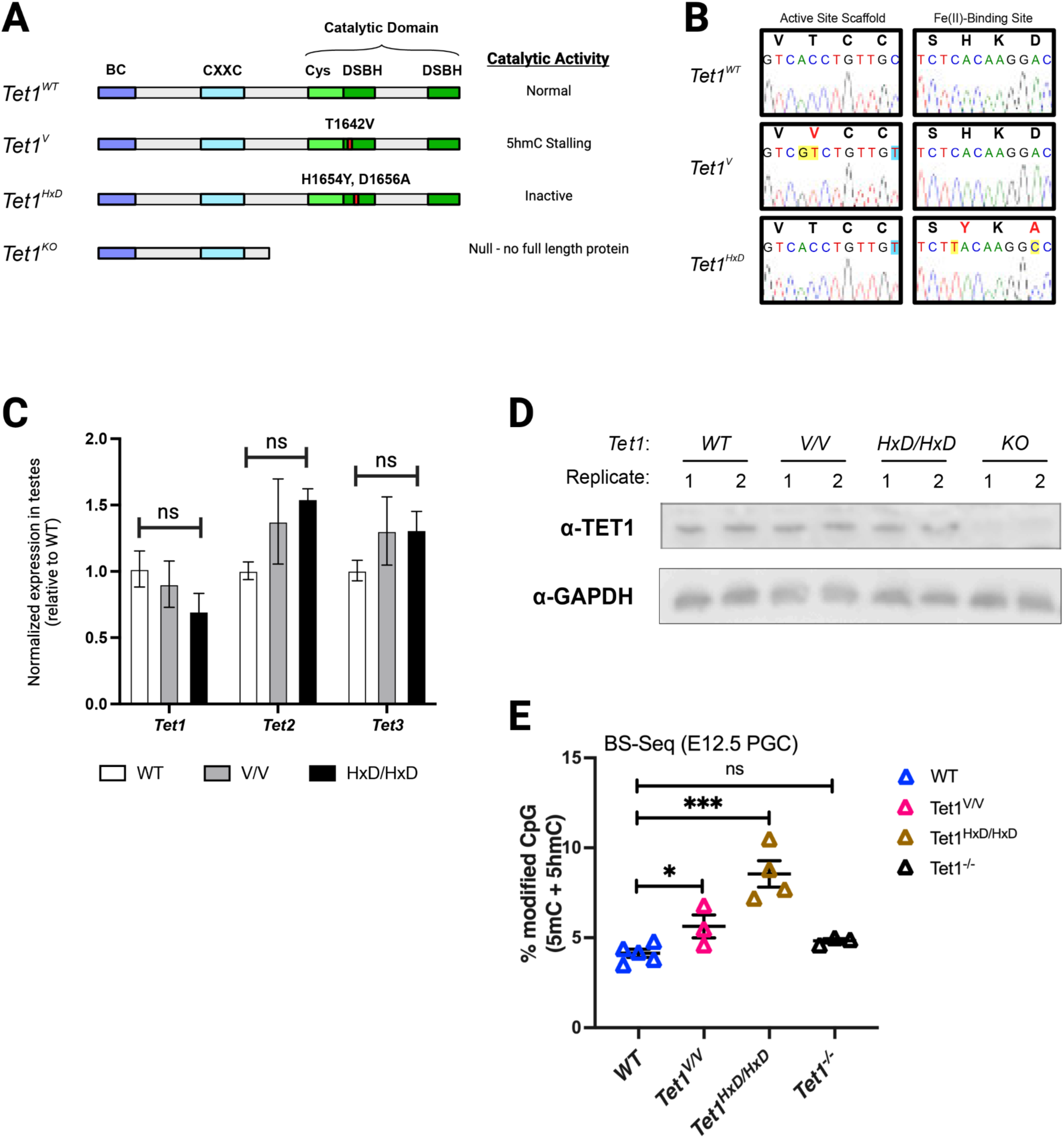
Generation and validation of 5hmC stalling *Tet1-V* and catalytically inactive *Tet1-HxD* mouse lines. A) Schematic representation of WT and mutant TET1 proteins. The N-terminal region of TET1 consists of the BC (“before CXXC”) and CXXC-type domains proposed to regulate DNA binding affinity and interactions with other regulatory factors. The C-terminal region consists of cysteine-rich (Cys) and double-stranded β-helix (DSBH) domains that comprise the catalytic domain. Threonine (T) to valine (V) substitution (T1642V) in the active site scaffold restricts further oxidation of 5hmC into 5fC and 5caC, resulting in the 5hmC stalling *Tet1-V* mouse line. Histidine (H) to tyrosine (Y) and aspartic acid (D) to alanine (A) substitutions (H1654Y, D1656A) abrogate Fe(II) cofactor binding resulting in the catalytically inactive form of TET1 (TET1-HxD). *Tet1^-/-^* line was generated previously and the absence of full-length or truncated protein was confirmed^78^. B) Sanger sequencing of *Tet1^+/+^*, *Tet1^V/V^*, and *Tet1^HxD/HxD^* alleles. C) Expression of *Tet1, Tet2*, and *Tet3* in testes of WT, *Tet1^V/V^* and *Tet1^HxD/HxD^* was measured by qRT-PCR (mean expression ± SEM; n=3, one-way ANOVA with Dunnett’s multiple comparisons test, normalized to *Nono* and *Rpl13*). D) Western blot for full length TET1 protein in testes of *Tet1^V/V^* and *Tet1^HxD/HxD^* animals with GAPDH as loading control. *Tet1^-/-^* samples are included as negative controls. E) Sparse BS-seq of *Tet1^+/+^*, *Tet1^V/V^*, and *Tet1^HxD/HxD^* E12.5 PGCs show global hypermethylation in *Tet1^V/V^* and *Tet1^HxD/HxD^* PGCs compared to WT. t-test vs WT, *p<0.05, ***p<0.0005.

Expression of *Tet1*, *Tet2*, and *Tet3* was measured in adult testis samples using quantitative real-time polymerase chain reaction (qRT-PCR). In homozygous *Tet1^V/V^* or *Tet1^HxD/HxD^* testes, *Tet1* expression was unchanged compared to wild type (WT). Catalytic mutant alleles also did not affect expression of *Tet2* or *Tet3* isoforms. We designed an RNA pyrosequencing assay to measure expression of the *Tet1^V^* and *Tet1^HxD^* alleles. Using cerebral cortex samples from heterozygous mice, where *Tet1* is actively transcribed, we determined that the mutant *Tet1^V^* or *Tet1^HxD^* alleles were expressed at equivalent levels to the *Tet1* WT allele (Supplemental figure 1F-G). Finally, full length TET1 protein can be detected in *Tet1^V/V^* and *Tet1^HxD/HxD^* testes at levels comparable to WT as assayed by Western blots (Figure 1D), indicating that proteins stability is unaffected in our *Tet1* mutants.

To evaluate global levels of 5mC in reprogramming PGCs, we employed sparse-coverage whole genome bisulfite sequencing (sparse BS-seq) to approximate 5mC and 5hmC levels^37, 38^. This method had previously been shown to accurately estimate global levels of genome methylation through sampling of at least 20,000 cytosines in the CpG context using next-generation sequencing and is amenable for low-input samples such as PGCs^38^. We benchmarked sparse BS-seq with liquid-chromatography-tandem mass spectrometry (LC-MS/MS) measurement of modified cytosines in the adult mouse cortex, a tissue where modified cytosines are relatively abundant (Supplemental Figure 2A)^39^. Our results showed good agreement between sparse-seq and LC-MS/MS in reporting global levels of modified 5mC (Supplemental Figure 2A).

In mice, germline epigenetic reprogramming occurs as PGCs migrate from the proximal epiblast to the bipotential gonad between embryonic day (E)7.25 and E13.5^40^. At E12.5 demethylation of the PGC genome is nearing completion^41^. Sparse-BS-seq of E12.5 PGCs revealed global hypermodification of *Tet1^V/V^* and *Tet1^HxD/HxD^* PGCs compared to WT, while *Tet1^-/-^*PGCs did not show elevated level of global modified cytosine, in agreement with previous reports (Figure 1E)^41–43^. These validations demonstrated that we generated two viable mouse*Tet1* catalytic mutants, which express full length TET1 proteins, with potentially distinct phenotypes from the previously established *Tet1^-/-^* lines^10, 44^. We proceeded to employ these mutants for delineation of catalytic and non-catalytic TET1 functions with respect to germline reprogramming.

### Catalytic Tet1 mutations lead to incomplete ICR reprogramming in sperm and imprinting defects in F1 offspring

ICRs are the most well-characterized loci to require TET1 during germline epigenetic reprogramming^10, 11^. Germ cells of male and female *Tet1^-/-^* mice show hypermethylation at ICRs, despite unchanged genome-wide methylation level^10, 11, 13^. However, while TET1 and 5hmC are detected in migrating PGCs, it is unknown whether 5hmC generation is sufficient to promote replication-coupled passive dilution or whether TET1-dependent ICR reprogramming requires the generation of 5fC/5caC^23^. To determine this, we measured the methylation of representative ICRs in sperm of *Tet1^V/V^*, *Tet1^HxD/HxD^*, and *Tet1^+/+^* mice (Figure 2A). *Peg1, Peg3, KvDMR*, and *Snrpn* are maternally methylated ICRs that are normally hypomethylated in sperm. *Tet1^V/V^* and *Tet1^HxD/HxD^* sperm showed *Peg1, Peg3*, and *KvDMR* hypermethylation compared to *Tet1^+/+^*, indicating incomplete ICR reprogramming in mutant germ cells (Figure 2A). Consistent with our previous report, *Snrpn* demethylation was not dependent on TET1, and its methylation was unaffected in *Tet1^V/V^* or *Tet1^HxD/HxD^* sperm^11^. The paternally methylated *H19/Igf2* ICR showed the expected hypermethylation in sperm for all genotypes.

**Figure 2.**
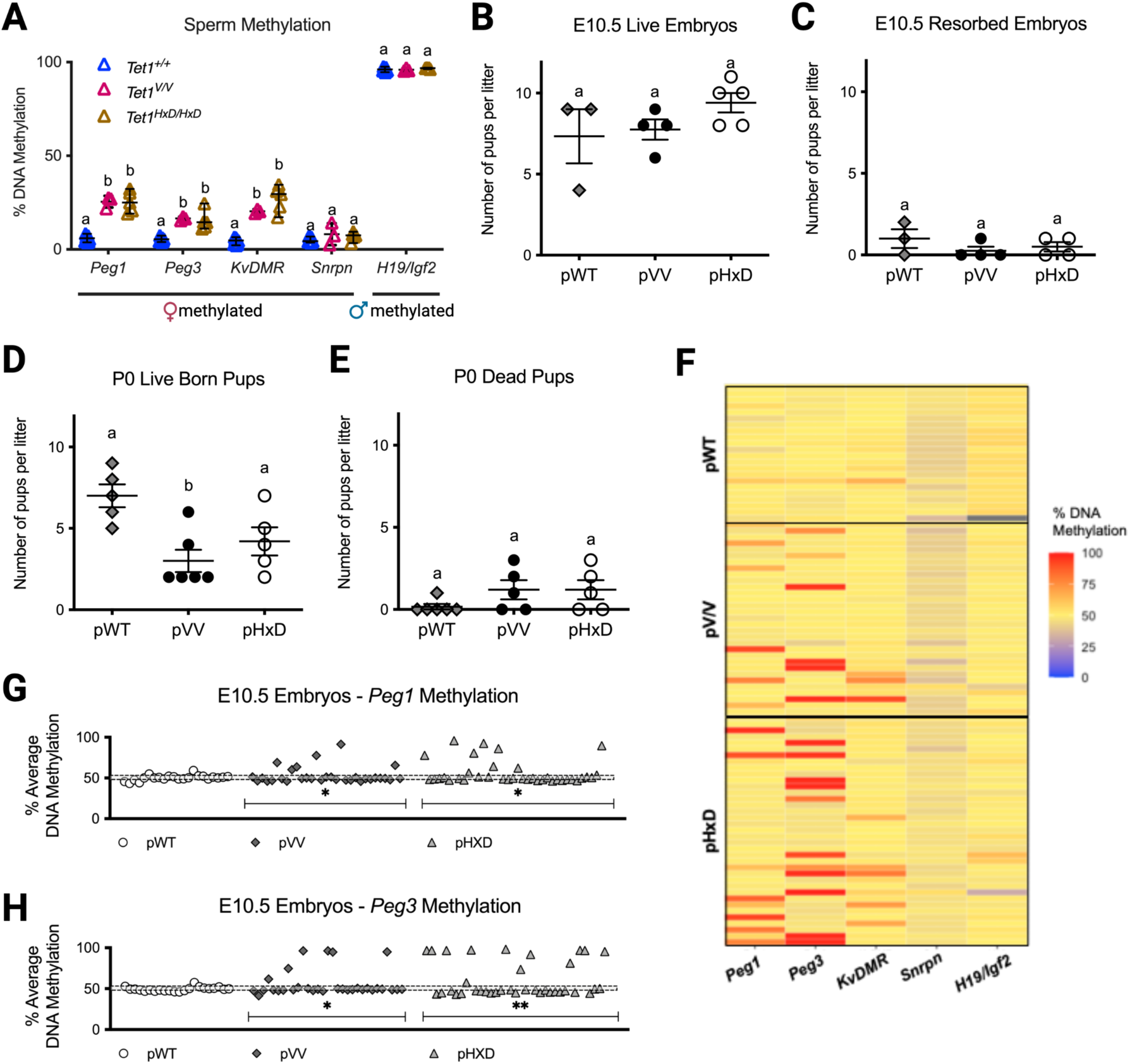
*Tet1^V/V^* and *Tet1^HxD/HxD^* males exhibit methylation defects at ICRs that are inherited by offspring. A) Methylation levels at maternally methylated ICRs *Peg1, Peg3*, *KvDMR*, and *Snrpn* as measured by pyrosequencing. Each data point represents methylation level of sperm sample from one adult mouse. *H19/Igf2* ICR is included as a paternally methylated ICR that exhibits full methylation in sperm (mean methylation ± SEM; n=3-5, one-way ANOVA with Dunnett’s multiple comparisons test, distinct letters indicate statistical significance). The number of live embryos (A) and resorbed embryos (B) per litter at E10.5 (mean number of pups per litter ± SEM, n=3-5 litters, one way ANOVA with Dunnett’s multiple comparisons test, distinct letters indicate statistical significance). The number of live pups (D) and dead pups (E) per litter at PND0 (mean number of pups per litter ± SEM, n=5-6 litters, one way ANOVA with Dunnett’s multiple comparisons test, distinct letters indicate statistical significance). F) Heatmap representation of DNA methylation levels at ICRs of all E10.5 offspring from *Tet1^+/+^*, *Tet1^V/V^* and *Tet1^HxD/HxD^* males. Each row represents an individual embryo of the indicated paternal genotype. The same offspring are depicted by ICR for *Peg1* (G) and *Peg3* (H) ICRs as measured by pyrosequencing. pWT n=22 embryos (3 litters), pVV n=31 embryos (4 litters), pHxD n=37 embryos (4 litters). *p<0.05, **p<0.01. Fisher’s exact test for frequency of hypermethylated embryo, shaded bars indicate average methylation of pWT embryos ± STDEV. Embryos with methylation above or below the shaded bar (+/-1 standard deviation of the mean of pWTs) are considered hyper- or hypomethylated and denoted as warmer color in the heatmap.

To test whether hypermethylated sperm of *Tet1* catalytic mutant males contributed to fertility, we mated *Tet1^V/V^*, *Tet1^HxD/HxD^*, or *Tet1^+/+^* males to C57BL6/J females (see Supplemental Figure 3 for breeding strategy). At midgestation (E10.5), pregnancies sired by *Tet1^V/V^* (pVV), *Tet1^HxD/HxD^* (pHxD) or WT (pWT) males showed equivalent numbers of developing (Figure 2B) and resorbed (Figure 2C) embryos. By birth (PND0), pVV showed significantly decreased litter size compared to pWT, while the decrease in pHxD litter size was not statistically significant (Figure 2D). Because we did not observe a significantly increased change in dead pups in pVV or pHxD litters at birth, we concluded that litter attrition likely occurred between E10.5 and birth (Figure 2E).

ICRs are protected from the post-fertilization global DNA demethylation that occurs in the embryo^45, 46^, and thus incomplete erasure of ICRs during germline development is expected to be stably inherited by TET1 mutant offspring (Supplemental Figure 3). Figure 2F depicts *Peg1*, *Peg3*, and *KvDMR* methylation levels as a heatmap at E10.5. *Snrpn* is included as maternally methylated ICR control that is not affected by TET1 mutations and *H19/Igf2* is included as paternally methylated ICR control. The number of hypermethylated offspring was significantly increased at *Peg1* and *Peg3* for pVV (19.35% for both) and pHxD (21.62% and 32.43%, respectively) compared to pWT (Figure 2G-H, Supplemental Figure 4A). Notably, the proportion of pVV and pHxD embryos exhibiting hypermethylation for a given ICR mirrored the degree of hypermethylation observed in *Tet1* mutant sperm. While most affected pVV or pHxD embryos only showed hypermethylation at one ICR, a few exceptions demonstrated hypermethylation at multiple ICRs (Figure 2F). However, no correlation was observed between DNA methylation levels at *Peg1, Peg3*, and *KvDMR* in individual embryos, suggesting independent segregation of alleles with affected loci during meiosis. We similarly measured hypermethylation incidence in pVV or pHxD PND0 brain and observed lower frequencies of affected pups compared to E10.5 embryos (Supplemental Figure 4C-H), consistent with ICR hypermethylation as a driver for increased embryonic lethality in *Tet1*-mutant offspring.

In summary, the hypermethylation of representative ICRs in *Tet1* catalytic mutant sperm and increased incidence of hypermethylated offspring of catalytic *Tet1* mutant males phenocopies that of *Tet1^-/-^* males that we previously reported^11^. Integrating across the variants, our findings demonstrate that, in contrast to the previously assumed role for 5hmC as a driver for replication-coupled passive dilution of 5mC, ICRs require TET1-mediated oxidation through 5fC and 5caC to achieve complete reprogramming in PGCs.

Global methylation analysis revealed partial rescue of Tet1^-/-^ sperm methylome by full length TET1^V^ and TET1^HxD^ catalytic mutant protein

During germline epigenetic reprogramming, DNA methylation is erased from the PGC genome to reset somatic methylation patterns. Previously, *Tet1* deletion was reported only to affect a relatively small number of late demethylating loci, which include ICRs and gametogenesis-related genes^10, 12, 13^. The methylome of *Tet1^-/-^* sperm DNA, however, had only been analyzed using reduced representative bisulfite sequencing, which limits the interrogation to regions enriched with CCGG motifs^10^. To determine the broader requirements for 5fC/5caC generation and TET1 non-catalytic activity, we employed Illumina’s Mouse Infinium Methylation BeadChip to assess the methylome of *Tet1^V/V,^ Tet1^HxD/HxD^*, *Tet1^-/-^* and WT sperm DNA. The BeadChip interrogates > 285,000 CpGs representative of the mouse genome with manually curated coverage of gene promoters, enhancers, repetitive elements, and known CpG islands, including regions relevant for imprinting biology^47^. We identified 1411 differentially methylated regions (DMRs, each DMR corresponds to a single probe on the array) in *Tet1^V/V^*, 2488 DMRs in *Tet1^HxD/HxD^*, and 6005 DMRs in *Tet1^-/-^* sperm compared to *Tet1^+/+^* (Figure 3A). While most DMRs in *Tet1^V/V^* and *Tet1^-/-^* sperm were hypermethylated (*Tet1^V/V^*: 1359 hypermethylated, 52 hypomethylated; *Tet1^-/-^*: 5631 hypermethylated, 374 hypomethylated), consistent with the role of TET1 in active demethylation, a substantial number of DMRs in *Tet1^HxD/HxD^* sperm were unexpectedly hypomethylated (1006 hypermethylated, 1482 hypomethylated). This result raises the possibility of a distinct function for the catalytically inactive TET1^HxD^ in DNA methylation, which may be unrelated to reprogramming.

**Figure 3.**
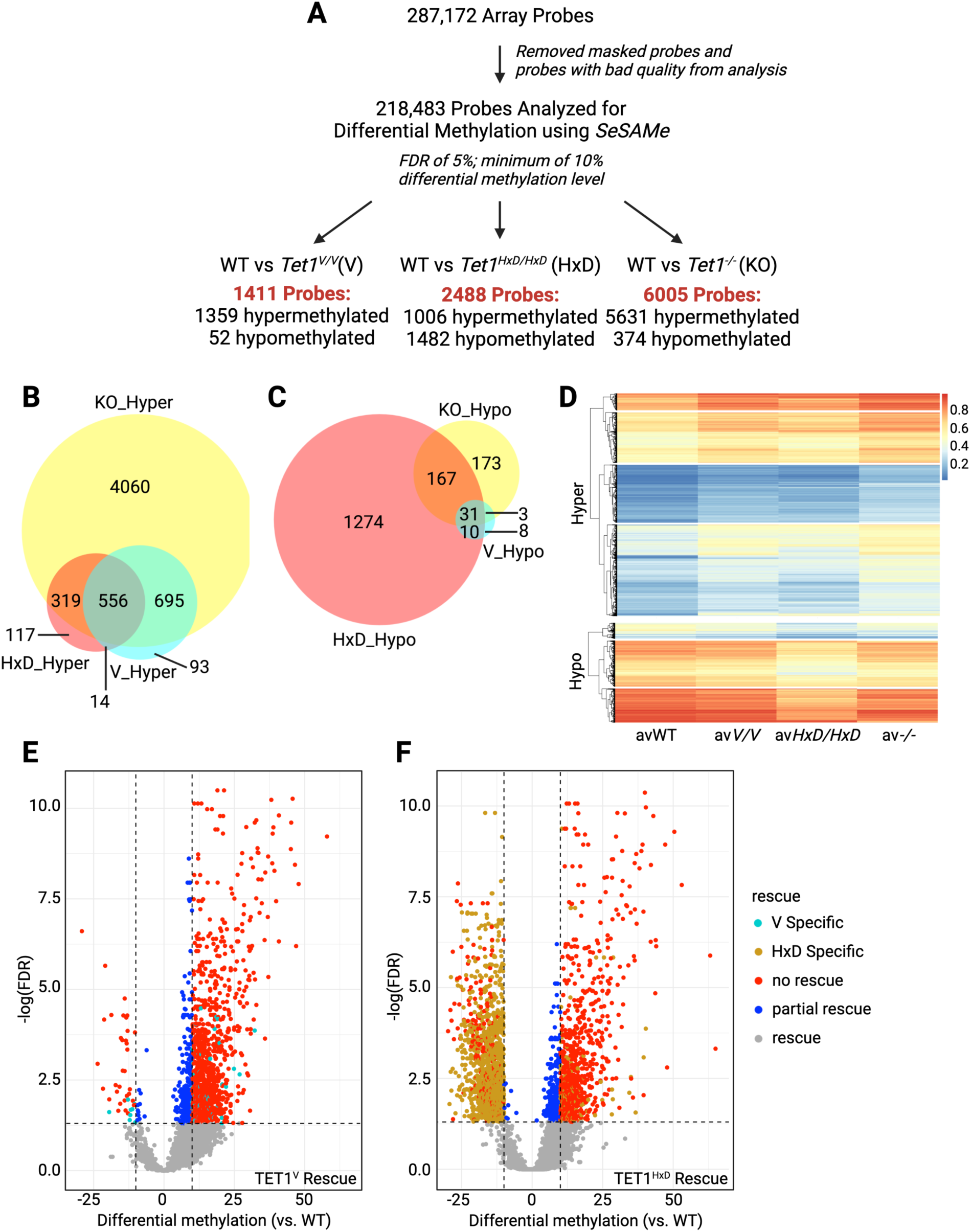
Global methylation analysis using Mouse Infinium Methylation BeadChip shows distinct methylome defects in catalytic mutant (*Tet1^V/V^, Tet1^HxD/HxD^*) sperm compared to *Tet1^-/-^* sperm. A) Flow chart showing differential methylation analysis of each *Tet1* mutant sperm sample compared to *Tet1^+/+^* (WT), n=8-10. A DMR is defined as a probe with FDR < 0.05 with minimum change in average methylation of greater than 10%. Venn overlap of significantly hypermethylated (B) and hypomethylated (C) DMRs in *Tet1^-/-^*, *Tet1^V/V^*, and *Tet1^HxD/HxD^* sperm compared to WT. D) Partially supervised clustering of methylation average for all hypermethylated DMRs (top) and all hypomethylated DMRs (bottom) identified in *Tet1^-/-^*, *Tet1^V/V^*, and *Tet1^HxD/HxD^* sperm. Note similar hypermethylated signatures of *Tet1^V/V^* and *Tet1^-/-^* sperm, and the distinct hypomethylated signature of *Tet1^HxD/HxD^* sperm. E) Volcano plot comparing the methylation status of *Tet1^-/-^* sperm DMRs to *Tet1^V/V^* sperm. F) Volcano plot comparing the methylation status of *Tet1^-/-^* sperm DMRs to *Tet1^HxD/HxD^* sperm. In these analyses, we assessed the methylation status of *Tet1^-/-^* DMRs within *Tet1^V/V^* or *Tet1^HxD/HxD^* samples. *Tet1^-/-^* DMRs that were no longer significant (FDR > 0.05) in *Tet1^V/V^* or *Tet1^HxD/HxD^* sperm were considered “rescue” (grey dots). *Tet1^-/-^* DMRs that remained significant (FDR < 0.05) but differential methylation average between mutant and WT <10% threshold were considered “partial rescue” (blue dots). “No rescue” (red dots) correspond to DMRs that were significant in both *Tet1^-/-^* and catalytic mutant sperm. DMRs that were changed only in *Tet1^V/V^* or *Tet1^HxD/HxD^* were plotted in the same volcano plots and denoted with aqua or yellow dots, respectively.

We next determined the overlap of hypermethylated and hypomethylated DMRs between the catalytic mutants (*Tet1^V/V^* and *Tet1^HxD/HxD^*) and *Tet1^-/-^* sperm (Figure 3B-C). 556 DMRs were commonly hypermethylated in all three mutants. These loci were classified as requiring TET1 catalytic activity to generate 5fC/5caC for complete reprogramming in the male germline (Figure 3B, Supplemental Table 1). Following NCBI reference sequence (RefSeq) annotation, we identified many imprinting-associated regions within these 556 DMRs (Supplemental Table 2), consistent with our previous conclusion that ICRs require the full competency of TET1 catalytic activity to achieve reprogramming (Figure 2). While the majority of hypermethylated DMRs in *Tet1^V/V^* and *Tet1^HxD/HxD^* overlapped *Tet1^-/-^* hypermethylated DMRs, *Tet1^HxD/HxD^* showed a large number of unique hypomethylated DMRs in sperm (Figure 3C). Partially supervised hierarchical clustering of DMR methylation levels clearly demonstrates the similarity of *Tet1^V/V^* and *Tet1^-/-^* hypermethylation signatures (Figure 3D, top) and the distinct hypomethylation signature of *Tet1^HxD/HxD^* (Figure 3D, bottom).

Significantly greater numbers of DMRs were identified in *Tet1^-/-^* sperm compared to either of the catalytic mutant sperm (6005 *Tet1^-/-^*vs 1411 *Tet1^V/V^* DMRs; 6005 *Tet1^-/-^* vs 2488 *Tet1^HxD/HxD^* DMRs), suggesting that TET1^V^ or TET1^HxD^ proteins can partially rescue the DNA methylation defects in the KO sperm. To clarify the degree of rescue that our new mutants provided, we assessed methylation levels and statistical significance of the 6005 DMRs of *Tet1^-/-^* sperm in *Tet1^V/V^* or *Tet1^HxD/HxD^* samples. *Tet1^-/-^* DMRs were considered rescued by full length TET1^V^ or TET^HxD^ if those DMRs no longer reached the threshold for statistical significance (FDR < 0.05), while partially rescued DMRs were still significantly altered in *Tet1^V/V^* or *Tet1^HxD/HxD^* samples but differential methylation between catalytic mutant samples and WT was no longer greater than 10%. Approximately 75% of KO DMRs were rescued by the expression of TET1^V^ (4369/6005, Figure 3E) or TET1^HxD^ (4679/6005, Figure 3F). The similar degrees of rescue that were demonstrated by the 5hmC dominant TET1^V^ and the catalytically inactive TET1^HxD^ suggest that majority of methylome defects observed in *Tet1^-/-^* sperm were due to the absence of TET1 non-catalytic activities, as full length TET1 with diminished or ablated 5mC oxidation activity is sufficient to achieve the WT methylation state at these loci. Alternatively, non-catalytic domains of TET1 protein may be sufficient to prevent aberrant *de novo* methylation at these loci during spermatogenesis. By contrast, only a modest number of KO DMRs were partially rescued in *Tet1^V/V^* and *Tet1^HxD/HxD^* samples (350 in *Tet1^V/V^* and 249 in *Tet1^HxD/HxD^*), perhaps representing a subset of loci where reprogramming is less efficient in the presence of TET1 catalytic mutants. Interestingly, these partially rescued KO DMRs are enriched at gene promoters, while fully rescued DMRs showed similar distribution to the totality of *Tet1^-/-^* DMRs (Supplemental Figure 5C). Finally, *Tet1^-/-^* DMRs that remained significantly altered in *Tet1^V/V^* (“no rescue”: 1286/6005) and *Tet1^HxD/HxD^* (“no rescue”: 1073/6005) sperm likely represent loci that are dependent on TET1’s catalytic activity to generate 5fC/5caC for reprogramming. Notably, with only 101 V-specific DMRs, the methylome defect of *Tet1^V/V^* sperm could be characterized as a less severe form of *Tet1^-/-^* sperm, while *Tet1^HxD/HxD^* exhibited a unique hypomethylation defect (1391 HxD specific DMRs). Taken together, the data indicate that the 5hmC-dominant TET1^V^ and catalytically inactive TET1^HxD^ rescue a significant proportion of the methylation defects observed in *Tet1^-/-^* sperm, thus supporting a more expansive role for TET1’s non-catalytic domains in male germline reprogramming.

### 5hmC-dominant Tet1^V/V^ and catalytically inactive Tet1^HxD/HxD^ sperm exhibit differential methylation in distinct genomic compartments

Consistent with previous reports that the loss of TET1 does not affect global levels of DNA methylation in PGCs or sperm, median methylation signals of *Tet1^V/V^*, *Tet1^HxD/HxD^*, and *Tet1^-/-^* sperm were not significantly different compared to WT in the Infinium Methylation BeadChip (Supplemental Figure 6A)^10, 13^. *Tet* loss of function mutants had previously been shown to exhibit hypermethylation of diverse genomic compartments including promoters, enhancers, and methylation canyons in the context of disease and development^32, 37, 48, 49^. To investigate the genomic context where methylation was most affected in *Tet1* mutant sperm, we annotated DMRs for genomic regions (Supplemental Figure 6B) and CpG density (Supplemental Figure 6C-E). For all three mutant genotypes, the largest proportion of DMRs mapped to intergenic regions (Supplemental Figure 6B). For *Tet1^V/V^* and *Tet1^-/-^*, which showed a similar hypermethylation signature (Figure 3D), the second most affected genomic compartments were exons, followed by introns. *Tet1^HxD/HxD^* sperm showed the largest proportion of DMRs in intergenic regions, followed by introns and exons. The lower proportion of DMRs mapping to promoter/transcriptional start site (TSS) regions is likely a function of promoters that are represented in the array^47^.

We used the R packages annotatr and AnnotationHub to determine the CpG densities where DMRs for each *Tet1* mutant mapped (Supplemental Figure 6C-E). These annotation packages define CpG shores as +/-2 kb from the ends of CpG islands and CpG shelves as +/-2 kb from the farthest limits of the CpG shores. Open sea CpG dinucleotides are located +/-4kb away from the end of a CpG island (Supplemental Figure 5B). The majority of hypermethylated DMRs were located in CpG sparse regions (“Open Sea”), followed by CpG islands. Hypomethylated DMRs that are dominant in *Tet1^HxD/HxD^* sperm mostly mapped to CpG sparse regions as well. The N-terminus of TET1 contains the CXXC domain that can bind to unmethylated CpGs^6, 50^. Because of this, it is thought that higher CpG density correlates with TET protein binding, and therefore greater reliance on TET function to maintain unmethylated state^6^. Our results demonstrate CpG density does not suggest TET1 dependence.

### Tet1-DMRs are associated with binding sites of methylation-sensitive and developmentally important transcription factors and chromatin states

To reveal the biological context of differential methylation that resulted from either loss of TET1 or expression of catalytic mutants TET1^V^ and TET1^HxD^ in the germline, we conducted feature enrichment analysis for each set of DMRs using knowYourCG (KYCG) function of the SeSAMe package^51^. Designed specifically for the Illumina methylation array, the KYCG algorithm considers that DMRs from the BeadChip array are a sparse representation of the methylome by clustering them by feature or motif enrichment. First, we assessed enrichment of hyper- or hypomethylated DMRs based on target biological design groups^47^ (Supplemental table 2). Hypermethylated DMRs for all three sperm genotypes showed the strongest enrichment for probes designed to target imprinting biology (KYCG term: ImprintDMR) and regions with known monoallelic methylation in adult tissues (KYCG term: MonoallelicMeth). Hypermethylated DMRs were also enriched for sperm unmethylated regions (KYCG term: SpermUnmeth). Of interest was the large number of hypomethylated DMRs in *Tet1^HxD/HxD^* sperm. The strongest probe design group enrichment for these hypomethylated DMRs, however, belongs to ∼60,000 randomly chosen CpGs with unknown biology that were included in the array for exploratory analysis (KYCG term: Random^47^), followed by predicted TSS of pseudogenes (KYCG term: pseudogenesTSS).

During PGC epigenetic reprogramming, DNA demethylation is tightly associated with chromatin remodeling that culminates in the activation of genes that are required for meiosis entry and continued gamete development^52–54^. While we conducted the global methylation analysis in *Tet1* mutant sperm, we hypothesize that the majority of observed methylation defects are the result of aberrant TET1 function during PGC reprogramming. Active DNA demethylation has been previously observed to be linked with transcription factor (TF) recruitment, especially at heterochromatic regions during reprogramming to pluripotency^20, 55–57^. We analyzed enrichment for TF binding motifs at hyper- and hypomethylated DMRs found in *Tet1^-/-^*, *Tet1^HxD/HxD^*, and *Tet1^V/V^* sperm (Figure 4A-C). For hypermethylated DMRs, binding motifs for MECP2 and DPPA2 were the most significantly enriched and overlapped with the largest number of DMRs in mutant sperm (left panels, Figure 4A-C). This result is consistent with the similar mechanisms of TET1, MECP2, and DPPA2 in targeting CpG dense regions of the genome that are particularly important during early embryonic development^6, 58–60^. Enrichment of TRIM28 and PRDM9 binding sites at hypermethylation DMRs was consistent with the roles of these TFs in meiosis and underscored TET1’s importance in maintaining normal methylation of regulatory regions central to germ cell development^12, 61–63^. In addition to TF binding motifs, we identified enrichment for targets of histone tail modifiers within hypermethylated DMRs. Target motifs of KDM4B, a H3K9 demethylase, are enriched in hypermethylated DMRs of all three *Tet* mutants, while motifs for KDM2A (H3K36 demethylase) and KDM2B (H3K4 and H3K36 demethylase) showed enrichment at *Tet1^-/-^* and *Tet1^HxD/HxD^* hypermethylated DMRs, respectively. This finding supports the hypothesis that active demethylation and histone reprogramming are tightly associated during germline reprogramming. Additionally, targeting motifs for SETDB1, an H3K9 methyltransferase, were enriched in hypermethylated DMRs of *Tet1^HxD/HxD^* and *Tet1^V/V^* sperm. Despite the substantial numbers of hypomethylated DMRs in *Tet1^HxD/HxD^* sperm, TF motif enrichment analysis was similarly uninformative as probe design group enrichment and only revealed weak association with CpG-associated methylation sensitive TFs such as CTCF, MBD1, MECP2, and DPPA2 (right panel, Figure 4B). Likewise, no enriched motifs were found for the 52 hypomethylated DMRs of *Tet1^V/V^* sperm.

**Figure 4.**
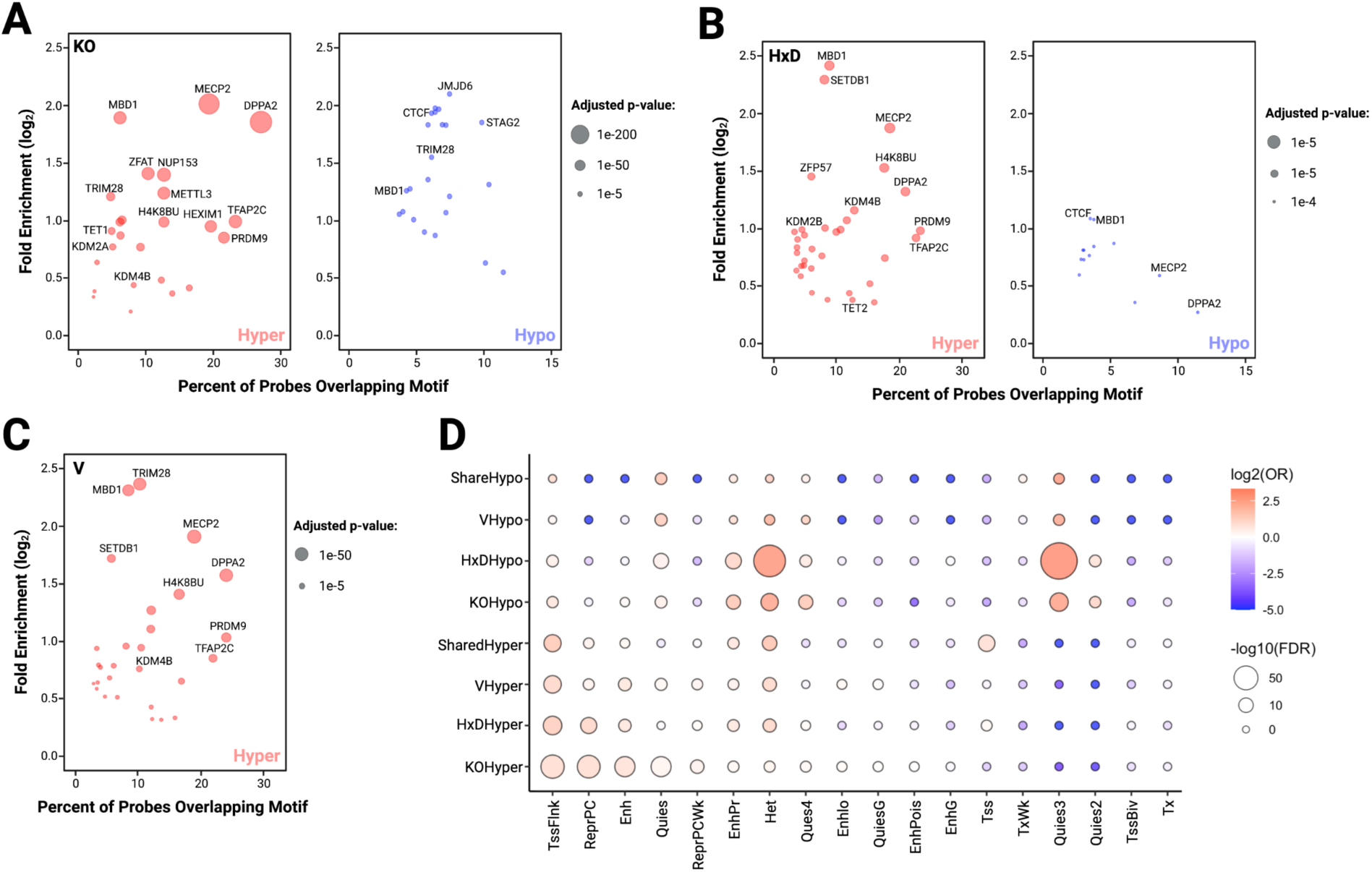
Transcription factors (TFs) and chromatin state enrichment at DMRs in *Tet1* mutant sperm. Transcription factors whose binding sites are enriched in hypermethylated DMRs (left) and in hypomethylated DMRs (right) for *Tet1^-/-^* (A), *Tet1^HxD/HxD^* (B), and *Tet1^V/V^* (C) sperm as identified by SeSAMe knowYourCG function. No transcription factor motif enrichment is found for hypomethylated DMRs in *Tet1^V/V^* sperm. Y-axis represents fold enrichment for the identified TF binding sites and X-axis represents the percentage of significant probes that overlap the binding sites. D) Enrichment DMRs in chromatin states as classified in ENCODE ChromHMM. Chromatin state enrichments are separated for hyper- and hypomethylated DMRs for each genotype.

Finally, we performed chromatin state discovery for DMRs using chromHMM (Figure 4D)^64^. Hypermethylated DMRs in *Tet1^-/-^*, *Tet1^V/V^*, and *Tet1^HxD/HxD^* as well as DMRs that are co-regulated among the three genotypes, showed strongest enrichment at TSS flanking region (TssFlnk), which may indicate non-genic, transcriptionally active regions such as enhancers^65^. Chromatin state discovery analysis was informative for hypomethylated DMRs that were specific to *Tet1^HxD/HxD^*, as these regions were enriched for heterochromatin chromatin state due to H3K9me3 localization (Het) or chromatin state that is associated with quiescence (Quies3)^66^. This finding suggests that expression of catalytically inactive TET1^HxD^ may cause derepression of silenced regions following methylation loss at these regions. Overall, these results support a scenario in which active demethylation and chromatin remodeling are likely to be closely coordinated during PGC reprogramming.

### Tet1 DMRs are located in regions that are excluded from de novo methylation in the hypermethylated sperm genome

In this study, we analyzed the sperm methylome that resulted from altered TET1 activities during PGC reprogramming. Upon the completion of epigenetic reprogramming, the male germline is hypermethylated late in gestation beginning around E15.5, which yields a highly methylated sperm genome compared to that of somatic cells or the oocyte genome^67, 68^. In the mouse, sperm genome global methylation is ∼90%, which is reflected in the median methylation signals of the array data (Supplemental Figure 6A)^68^. While *de novo* methylation occurs indiscriminately in prospermatogonia, there are regions that are excluded from the action of *de novo* DNMT DNMT3A/3L and thereby protected from gaining methylation in the sperm genome^36, 69^. Among these sperm-specific hypomethylated regions (HMRs) are ICRs that are methylated only on the maternal allele (Figure 5B)^70^. KYCG analysis indicated that DMRs found in mutant sperm are enriched for probes that target imprinting biology (Supplemental Table 2). Using Bernoulli’s distribution testing, we determined that imprinting probes, which make up 0.2% of the array, were overrepresented in the *Tet1* mutant DMRs, particularly in *Tet1^V/V^* sperm (Figure 5A).

**Figure 5.**
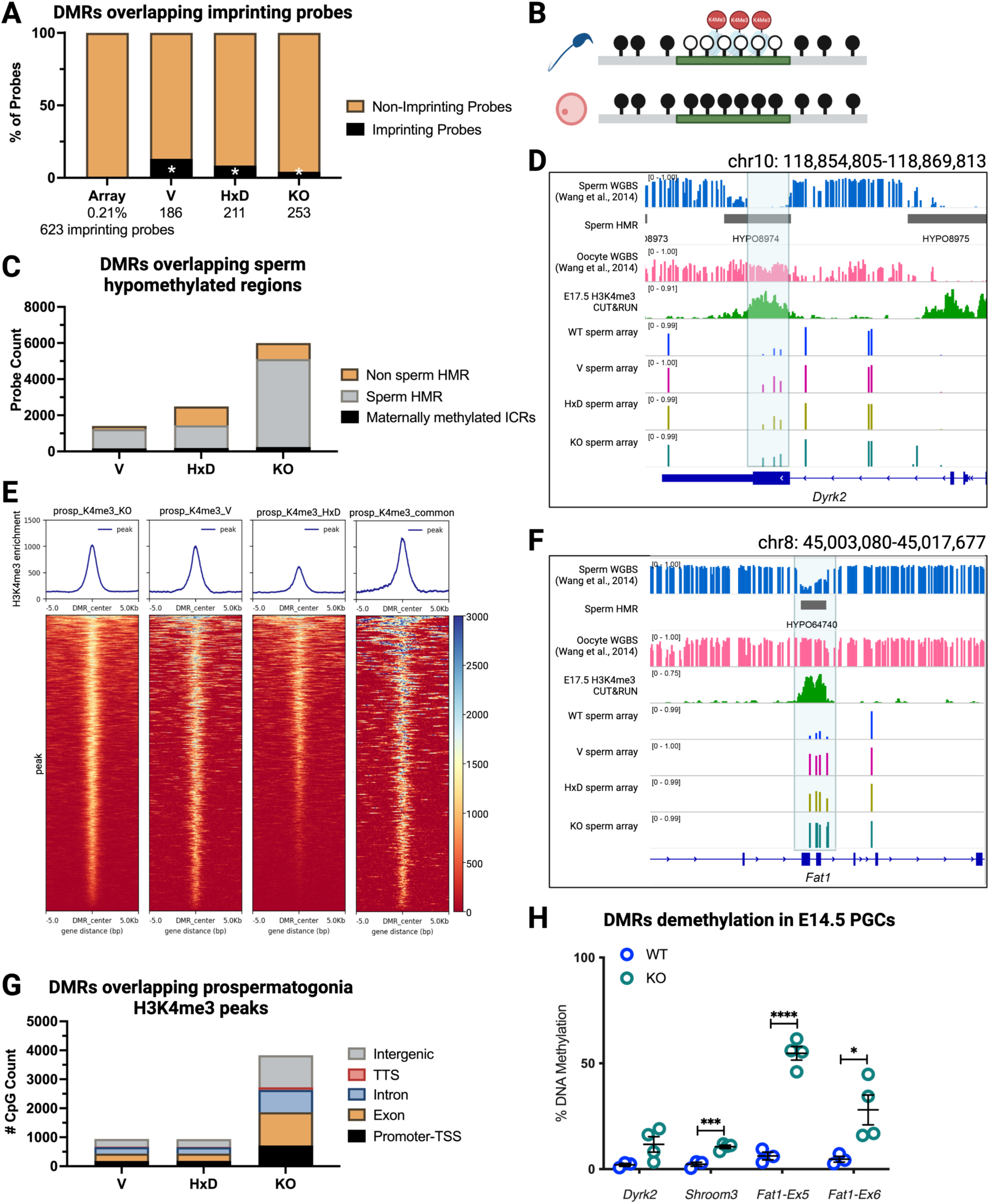
Identification of TET1-dependent sperm-specific hypomethylated regions. A) Distribution of DMRs classified as related to imprinting biology in the array annotation. *p-value < 0.05; two-sided Bernoulli distribution test as compared to all probes in the array. B) Cartoon of a sperm hypomethylated region that is excluded from *de novo* methylation through enrichment of H3K4me3. C) Distribution of DMRs that overlap unmethylated regions in the sperm genome (sperm HMRs) as determined by DNMTools function “hmr”, which identified methylation canyons within a WGBS dataset (GEO: GSE56697^72^). D, F) Representative examples of regions that are commonly hypermethylated in *Tet1^-/-^*, *Tet1^V/V^, Tet1^HxD/HxD^* sperm with overlap to sperm hypomethylated regions and H3K4me3 enrichment during *de novo* methylation in E17.5 prospermatogonia. Sperm HMRs are indicated with grey bars. E) Heatmaps and metaplots of E17.5 prospermatogonia H3K4me3 enrichment centered on DMRs for each genotype or those that are shared among three mutants, measured in counts per million (CPM). G) Genomic distribution of DMRs that overlap with H3K4me3 enrichment in E17.5 prospermatogonia as annotated by HOMER. H) Methylation analysis of newly identified TET1-dependent sperm hypomethylated regions in demethylating WT and *Tet1^-/-^* E14.5 PGCs using targeted bisulfite sequencing. n=3-4, *p-value < 0.05, ***p-value < 0.001, ****p-value < 0.0001, two-tail t-test.

We then asked whether DMRs of *Tet1* mutant sperm are located in sperm-specific HMRs unrelated to imprinting. In particular, we were interested in hypermethylated DMRs, which would be consistent with regions that utilize TET1 for active demethylation. We identified sperm-specific HMRs using DNMTools HMR algorithm, which searches for methylation canyons in WGBS datasets^71^. Using previously published sperm WGBS (GEO: GSE56697^72^), DNMTools discovered 76,227 distinct sperm HMRs, which we overlapped with DMRs for each *Tet1* mutant genotype (Figure 5C, Supplemental Table 3). A substantial portion of DMRs in *Tet1^V/V^* (87%, 1235/1411), *Tet1^HxD/HxD^* (58%; 1444/2488), and *Tet1^-/-^* sperm (85%; 5113/6005) were indeed located within sperm-specific HMRs. The lower proportion of *Tet1^HxD/HxD^* DMRs within sperm HMRs compared to *Tet1^V/V^* and *Tet1^-/-^* is a function of the hypomethylated DMRs, which are specific to the catalytically inactive genotype and fully excluded from HMRs. We also assessed whether DMRs fell within sperm HMRs that are typically methylated in the oocyte (Figure 5B, such as ICRs), and confirmed that, in addition to ICRs, 25-40% of DMRs, depending on the genotype, were located in regions that are differentially methylated between the sperm and the oocyte genomes (Supplemental Figure 7A).

As stated above, *de novo* methylation in the male germline results in a highly methylated sperm genome. The DNMT3A complex, however, is inhibited by H3K4me3-enriched regions^67^. We performed CUT&RUN for H3K4me3 on WT prospermatogonia to identify regions that are enriched for this chromatin mark. Genome-wide, H3K4me3 signals of E17.5 prospermatogonia showed strong overlap with the sperm HMRs, suggesting that methylation was indeed excluded from regions that are enriched for H3K4me3 (Supplemental Figure 7B). We next overlapped prospermatogonia H3K4me3 signals with *Tet1* mutant DMRs to determine 1) if DMRs were enriched in regions typically excluded from *de novo* methylation, and 2) whether dysregulated DNA methylation occurred in regulatory regions. We observed a significant presence of H3K4me3 signals corresponding to *Tet1^-/-^*, *Tet1^V/V^*, and *Tet1^HxD/HxD^* DMRs (Figure 6E, Supplemental Table 3). Overall, 3841/6005 *Tet1^-/-^*, 942/1411 *Tet1^V/V^*, and 939/2488 *Tet1^HxD/HxD^*DMRs overlapped with H3K4me3 peaks in prospermatogonia. Because H3K4me3 is more commonly used as a marker of active or bivalent promoters, we assessed the genomic locations of H3K4me3-overlapping DMRs. Counter to our expectations, most DMRs that overlapped with H3K4me3 regions were located within gene bodies (exons and introns) or intergenic regions instead of annotated promoters or TSSs (Figure 5G). Indeed, only ∼20% of DMRs that overlapped with H3K4me3 peaks were mapped to annotated promoters, regardless of the *Tet1* genotype (Figure 5G). To determine whether these DMRs mapped to defined promoters at later stages of spermatogenesis, we overlapped *Tet1* mutant DMRs with previously published H3K4me3 ChIP-seq data sets for spermatogonia^35^, pachytene spermatocyte^73^, round spermatid^73^, and sperm^74^ (Supplemental Figure 7E-G) and found that the *Tet1* mutant, H3K4me3-overlapping DMRs were not located within gene promoters. Representative examples of DMRs in non-promoter H3K4me3-marked regions include exon 3 of *Dyrk2* and exon5-6 of *Fat1* (Figure 5D, F). While these regions are expected to repel DNMT3 action during *de novo* methylation to generate sperm HMRs during normal germline development, they are hypermethylated in *Tet1^V/V^*, *Tet1^HxD/HxD^*, and *Tet1^-/-^* sperm, supporting the requirement of TET1 catalytic activity to help curtail ectopic 5mC deposition during germline reprogramming. Although these H3K4me3 peaks did not fall within gene promoters, we noted that similar to ICRs (*Peg1* is shown here as example, Supplemental Figure 7D), H3K4me3 enrichment remained throughout spermatogenesis at these *Tet1* mutant DMRs (e.g. *Fat1*, Supplemental Figure 7B), even in post meiotic spermatids and sperm where the majority of histones are replaced by protamines (Supplemental Figure 7E-G). Finally, we assessed methylation levels of select non-ICR, TET1-dependent sperm HMRs in E14.5 WT and *Tet1^-/-^* PGCs, a time point that marks the completion of germline reprogramming. Methylation analysis confirmed that TET1-dependent sperm HMRs failed to demethylate completely at this stage (Figure 5H). Taken together, our data demonstrated a dependency for active DNA demethylation during reprogramming in regions that will eventually be excluded from methylation in the male germline.

**Figure 6.**
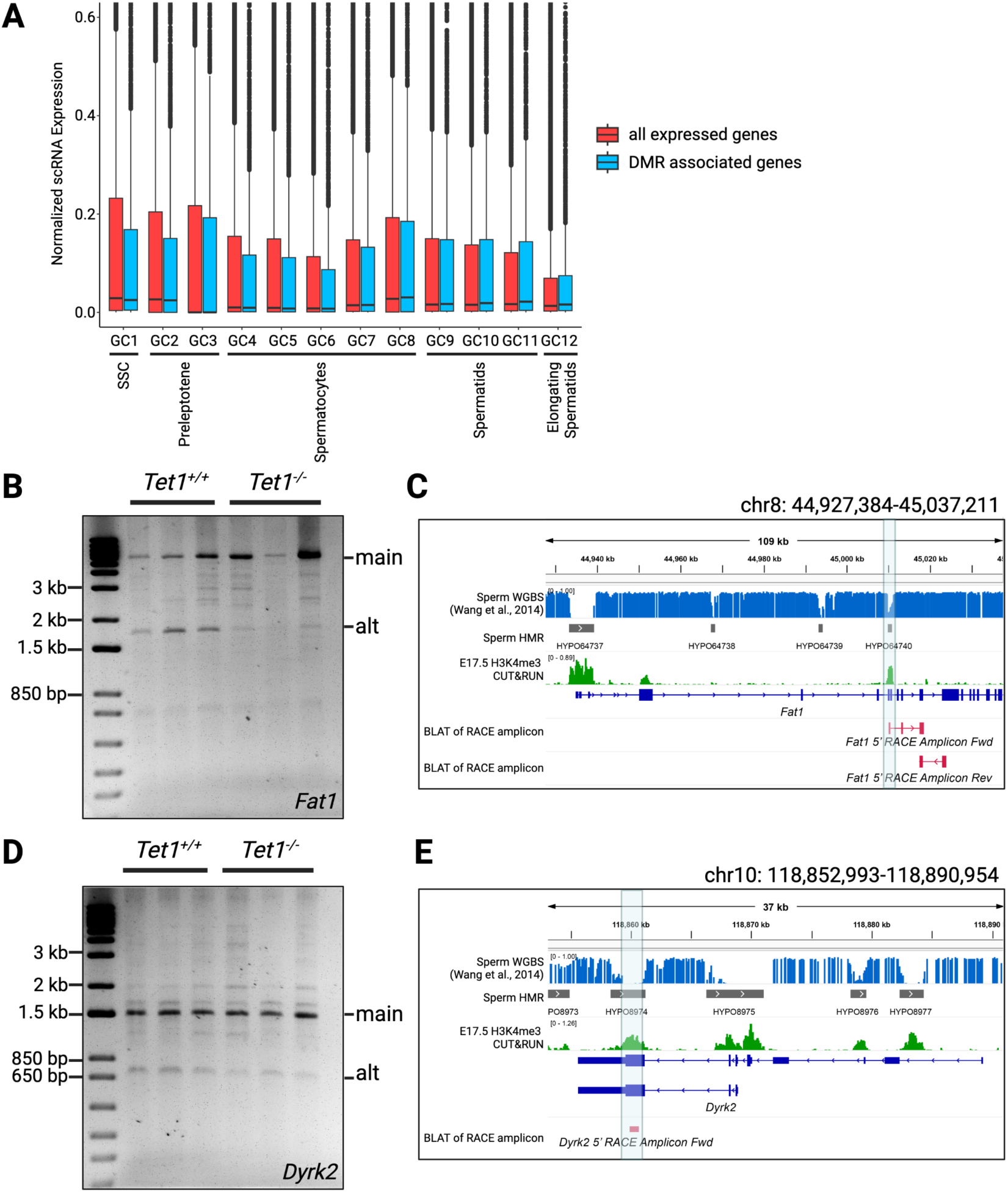
DMR-associated genes are expressed throughout spermatogenesis. A) Comparison of DMR-associated gene expression and all genes expressed throughout spermatogenesis based on normalized gene expression from publicly available scRNAseq (GEO: GSE112393^75^). B) *Fat1* 5’ RACE of *Tet1^+/+^* and *Tet1^-/-^* testes cDNA where the alternative product (alt) at 1586 kb is detected in addition to the main product (main) at ∼5.5 kb. C) BLAT of *Fat1* RACE alternative amplicon mapped to *Fat1* DMR. D) *Dyrk2* 5’ RACE of *Tet1^+/+^* and *Tet1^-/-^* testes cDNA where the main product is ∼1.5 kb and alternate is 745 bp. E) BLAT of *Dyrk2* RACE alternative amplicon mapped to *Dyrk2* DMR.

### Genes associated with Tet1 DMRs are expressed throughout spermatogenesis

We referred to publicly available single-cell RNA sequencing (scRNAseq) data of adult mouse testes to determine the spermatogenesis stages in which genes associated with TET1-dependent DMRs are expressed^75^. Previous analysis of the scRNAseq data identified 12 germ cell clusters that correspond to all developmental stages found in adult testes, including spermatogonial stem cells (SSC, Figure 6A, GC1), two stages of transitional preleptotene (GC2-3), 5 stages of spermatocytes undergoing meiosis (GC4-8), 3 stages of post meiotic spermatids (GC9-11), and elongating spermatids (GC12)^75^. We mapped all DMRs to the nearest annotated genes, which resulted in 4358 DMR-associated genes. 3207 of these genes have detectable expression in at least one GC stage within the scRNAseq data set (Figure 6A). DMR-associated genes are expressed in all germ cell clusters, with median expression level comparable to that of all expressed genes at each stage of spermatogenesis (Figure 6A).

A subset of DMRs overlapped with H3K4me3 peaks outside of known promoters throughout all analyzed germ cell stages (Supplemental Figure 7B, E-G). We next investigated whether these regions may serve as alternative TSSs. We conducted rapid amplification of cDNA ends (RACE) to identify alternative 5’ ends on the *Fat1* and *Dyrk2* loci (Figure 6B-E) using adult testes cDNA to obtain transcript pools from all spermatogenic stages. From the scRNAseq data, *Fat1* is expressed in GC1 cluster (SSC), while *Dyrk2* is expressed in GC8-GC11 clusters (late spermatocyte, spermatids). Using 3’ gene specific primers downstream of the DMRs, we detected alternative amplicons (Figure 6B, D) that are shorter than the expected transcripts from the RefSeq annotated promoters at each locus (*Fat1* main: ∼5.5 kb, *Fat1* alt: 1586 kb; *Dyrk2* main: ∼1.5 kb, *Dyrk2* alt: 745 bp). We subcloned and sequenced the alternative amplicons for *Fat1 and Dyrk2*, which mapped their 5’ end to the DMRs at exon 5-6 of *Fat1* and exon 3 of *Dyrk2* (Figure 6C, E). These results demonstrated that at select loci, TET1-dependent sperm hypomethylated regions can be used as alternative transcriptional start sites during spermatogenesis.

## Discussion

The discovery of TET proteins as DNA dioxygenases led to a paradigm shift in epigenetics field, where it was previously thought that DNA methylation was largely erased through the lack of maintenance during cell division. There is now ample evidence for the importance of active DNA demethylation pathways during reprogramming to pluripotency and during PGC epigenetic reprogramming^10–13, 37, 76–79^. What remains to be answered are questions such as the non-catalytic functions of TET as epigenetic regulators and the regulatory potential of higher order oxidized cytosine bases (i.e. 5fC and 5caC) within the genome. Expanding upon the biochemical discovery of functional mutations within the catalytic domain of TET1 and TET2 proteins that alter the enzymes’ catalytic activities, we generated 5hmC dominant *Tet1^V^* and catalytically inactive *Tet1^HxD^* mouse lines to 1) study the importance of 5fC/5caC generation, and 2) elucidate the function of TET1’s non-catalytic regulatory activity in the context of PGC epigenetic reprogramming^34, 37^.

Our results show that ICRs, the most well characterized TET1-dependent loci during germline reprogramming, require 5fC/5caC generation to achieve complete reprogramming in the male germline, as evidenced by *Tet1^V/V^* males exhibiting similar heritable imprinting defects in sperm as *Tet1^-/-^*mice^11^. This finding contradicts previous views that the role of TET in germline reprogramming is restricted to the generation of 5hmC to repel DNMT1 binding and promote passive dilution^13, 23, 24, 26^. We also conducted genome-wide methylation analysis using the Infinium Methylation BeadChip to assess methylation patterns in WT, *Tet1^V/V^*, *Tet1^HxD/HxD^*, and *Tet1^-/-^* sperm. This allowed us to analyze the methylomes of numerous samples at base-resolution for ∼280,000 representative CpGs across the mouse genome and revealed more extensive methylome defects than previously reported in *Tet1^-/-^* sperm^10, 13^.

Comparison to the *Tet1^-/-^* sperm methylome is essential in our studies because it reveals the distinct methylation defects that result from the presence of full length 5hmC-dominant TET1^V^ or the catalytically inactive TET1^HxD^ proteins (Figure 3). Of note, we showed that TET1^V^ and TET1^HxD^ can partially rescue the hypermethylation phenotype of *Tet1^-/-^* sperm. Interestingly, 5hmC-dominant TET1^V^ and catalytically inactive TET1^HxD^ rescue *Tet1^-/-^* hypermethylated loci to a similar degree (Figure 3E, F), suggesting that a large proportion of TET1-dependent loci in the germline do not actually require its catalytic processivity to achieve or maintain normal methylation. TET1 contains a CXXC zinc finger domain within its N-terminus that specifically recognizes unmethylated CpG in clusters, although it is unknown whether TET1 can act as a safeguard against *de novo* methylation by acting as a physical hindrance for DNMTs while binding to unmethylated CpGs^80, 81^. It is plausible that DMRs that are rescued by the presence of full length, catalytic mutant forms of TET1 are those that do not require TET1 for active demethylation, but instead require protection from the DNMT3 complex during *de novo* methylation prospermatogonia, either through TET1 physical localization or through the contribution of TET1 in recruiting histone modifiers to generate a DNMT3 repelling chromatin environment.

The role of TET1 in patterning the sperm methylome is supported by the finding that the majority of DMRs in all three mutant genotypes overlap sperm-specific hypomethylated regions (HMRs). The sperm genome is uniquely hypermethylated; this methylation pattern is acquired during the prospermatogonia stage when DNMT3A/3L complex targets the full genome, excluding protected regions^35, 67, 82, 83^. To date, H3K4me3 is thought to be the most dominant DNMT3-repelling epigenetic mark in the germline genome^67^. We determined that *Tet1*-mutant DMRs overlapped with regions that are enriched for H3K4me3 in prospermatogonia, suggesting these regions are normally excluded from DNA methylation. This result is similar to previous reports in mouse embryonic fibroblasts, where combined loss of TET1 and TET2 promoted methylation invasion into discrete hypomethylated canyons within the genome^84^. It is not entirely understood why the hypermethylated sperm genome is interspersed with discrete HMRs. Included within these regions are maternally methylated ICRs, as well as evolutionarily conserved TSS and subfamilies of retrotransposable elements that are species specific^3, 36, 69^. Clinically, hypermethylation of sperm HMRs has been associated with idiopathic infertility and poor outcomes in couples undergoing fertility treatments^85–88^. We confirmed the TET1 requirement for reprogramming in representative non-ICR sperm HMRs, suggesting that in the sperm genome, regions fated for methylation-exclusion may originate as late-demethylating loci that require TET1-mediated active demethylation. TET1 may also contribute to the formation of a DNMT3-repelling chromatin environment at these loci through interaction with OGT to direct the activity of COMPASS family H3K4 methyltransferase^28^.

Interestingly, the majority of TET1-dependent, H3K4me3-enriched sperm HMRs are located within gene bodies rather than annotated promoters. Using 5’ RACE, we identified Tet1-DMRs at gene bodies of *Fat1* and *Dyrk2* as bona fide alternative promoters. FAT1 is a protocadherin that controls cell proliferation. Mice deficient for *Fat1* exhibit perinatal lethality due to defects in adhesion junctions of regal glomerular epithelial cells^89^. DYRK2 is a dual specificity tyrosine kinase that regulates ciliogenesis and Hedgehog signaling during embryogenesis^90^. While these genes have not been previously studied in spermatogenesis, scRNAseq data confirmed their expression in germ cell development. Surprisingly, while abundant expression of lincRNAs and alternatively spliced RNAs has been described for the testes, the use of alternative promoters during spermatogenesis has not been studied in mammals^91, 92^. In *Drosophila*, the transition from proliferating spermatogonia to differentiating spermatocytes is accompanied by dramatic change in transcription, with about a third of new transcripts originating from alternative promoters^93^. Some uses of alternative promoters include rendering tissue specificity, regulating timing of expression during development, and controlling translation efficiency of transcripts^94^. While the function of mammalian sperm HMRs remains enigmatic and underexplored, our study reveals at least one compelling use of these regions as alternative transcriptional start sites during spermatogenesis.

The association between TET1 and chromatin remodeling is further supported in our study by the enrichment of binding sites for multiple histone modifiers at *Tet1* mutant DMRs (Figure 4). These results are consistent with extensive chromatin remodeling that is concurrent with DNA methylation erasure during germline reprogramming, including the depletion of H3K9me2 and redistribution of H3K27me3 and H3K9ac in the PGC genome^52, 95^. In ESCs, TET1, PRC2 (an H3K27me3 writer), and the deacetylase complex Sin3a co-occupy similar genomic regions^32^. While *Tet1* deficiency causes PRC2 loss and Sin3a enrichment at bivalent promoters in ESCs, expression of catalytically inactive *Tet1^m/m^* (similar to our *Tet1^HxD^* allele) does not affect the localization of these histone modifiers, suggesting that TET1 is acting through its non-catalytic domains at such loci^32^. The two mouse models developed in this study will provide additional opportunities to explore the relationship between chromatin remodeling and TET1 non-catalytic functions.

One of the unexpected findings in catalytically inactive *Tet1^HxD/HxD^* sperm is the unique hypomethylation signature that is not observed in other mutants. There is currently scant data on the relative affinity of TET’s catalytic domain for oxidized cytosine bases compared to 5mC or unmethylated cytosine. If the TET catalytic domain prefers 5mC over 5hmC, it is possible that loss of 5hmC catalysis in *Tet1^HxD/HxD^* sperm may promote prolonged occupancy by TET1^HxD^, which then mediates the recruitment of chromatin modifiers to form a DNMT-inaccessible environment. Notably, the loss of methylation in *Tet1^HxD/HxD^* sperm occurs in heterochromatic regions, further suggesting that methylation dilution is likely to be accompanied by a change in the chromatin state. Figure 7 summarizes our proposed model of differences in TET1^V^ and TET1^HxD^ action during PGC reprogramming, as well as the subsequent consequences, including changes to the chromatin environment and physical occupancy on the genome, during *de novo* methylation.

**Figure 7.**
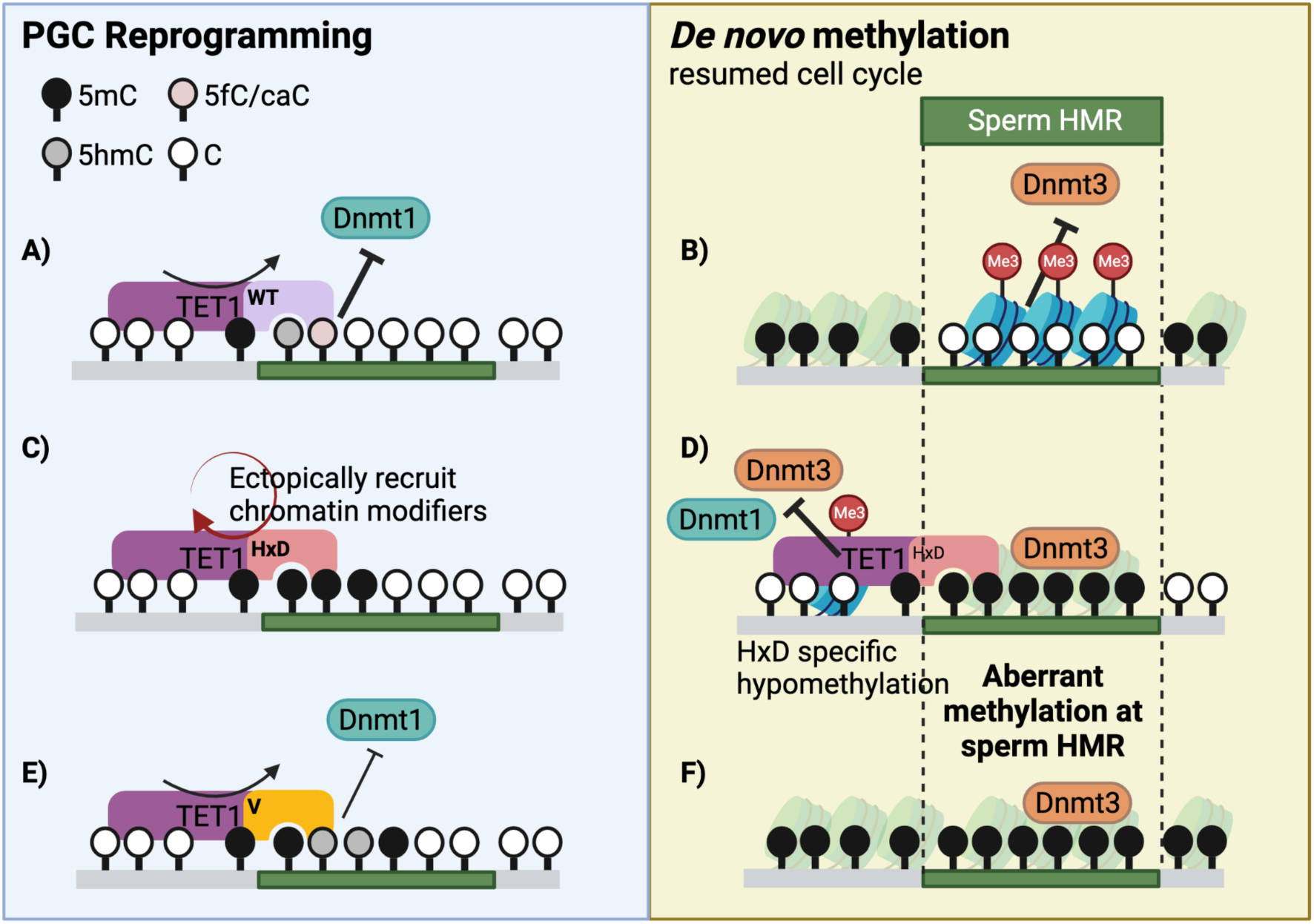
Proposed model of TET1-catalytic mutant access during germline reprogramming. TET1 generation of 5hmC and higher ordered oxidized bases 5fC/5caC is required for demethylation of sperm HMRs, which are generated via exclusion of DNMT3 during *de novo* methylation in prospermatogonia (A-B). Catalytically inactive TET1^HxD^ may cause prolonged occupancy at TET1-dependent loci, leading to ectopic recruitment of chromatin modifiers - curved arrows indicating release of the locus following catalysis in TET1^WT^ or TET1^V^ (C). In prospermatogonia, DNMT3 accesses sperm HMRs, creating hypermethylated regions (D). Physical hindrance by extended TET1^HxD^occupancy or aberrant chromatin environment causes HxD-specific hypomethylated DMRs. Similar to TET1^WT^, generation of 5hmC by TET1^V^ allows for TET1 turnover. However, 5hmC generation is not sufficient to maintain hypomethylated states at sperm HMRs (E-F).

In summary, our work demonstrates distinct methylation defects in the male germline following expression of 5hmC stalling TET1*^V^* or the catalytically inactive TET1^HxD^ mutants compared to *Tet1^-/-^*, allowing us to determine the dependency of select loci on 5fC/5caC generation to achieve complete reprogramming. These 5fC/5caC-reliant loci encompass a larger portion of the male germline genome than the previously thought. In addition to ICRs, we identified numerous sperm hypomethylated regions dependent on TET1 for either reprogramming or protection from *de novo* methylation. Overall, our study supports the roles of TET1 not only as a component of germline reprogramming, but also as a contributor to patterning the eventual sperm epigenome by influencing *de novo* methylation in prospermatogonia.

## Limitations of study

We employed Illumina Mouse Infinium BeadChip array for our whole genome analysis of the sperm methylome. While the array was designed to be a good representation of the mouse genome, only a minority of CpGs within the genome (∼285,000 CpGs) are examined. There are many more relevant CpGs, including those at ICRs, where probe design is not feasible. Moreover, the array is overrepresented for cancer- and aging-related promoters and CpGs at open sea (sparse regions). WGBS will likely capture many more TET1-dependent regions as well as regions that are uniquely altered in *Tet1^V/V^*, *Tet1^HxD/Hx^*^D^, and *Tet1^-/-^* sperm.

## Acknowledgements

This work was supported by National Institute of Health grant numbers R01GM146388 (MSB), R01GM051279 (MSB), R01GM118501 (RMK), R01HG010646 (RMK), F32HD101230 (RDP), and F31HD098764 (BAC). We thank Yemin Lan for advice with bioinformatics. We acknowledge the Children’s Hospital of Philadelphia Flow Cytometry Core and the Center for Applied Genomics for performing the Infinium Mouse Methylation BeadChip assays.

## Author Contributions

Experiments were designed by RDP and BAC with assistance and supervision from MSB and RMK. BAC and NAL developed the *Tet1^V^* and *Tet1^HxD^* allele CRISPR/Cas9 mutagenesis strategy. RDP performed and analyzed most experiments with the following exceptions: BAC designed and generated the CRISPR/Cas9 strategy for mutant mice initially characterized lines (Sanger sequencing, Western blot, qRT-PCR), SAC performed the CUT&RUN experiment, SW optimized 5’-RACE, JMF performed sparse-seq validation on LC-MS/MS, RDP, BAC, and ZL conducted offspring methylation assays. Global methylation data analysis was performed by RDP and BAC, CUT&RUN data analysis was performed by RDP, and ChIP-seq data analysis performed by ZL and RDP. RDP wrote the manuscript, and the manuscript was approved by all authors.

---

**Supplemental Figure 1.**
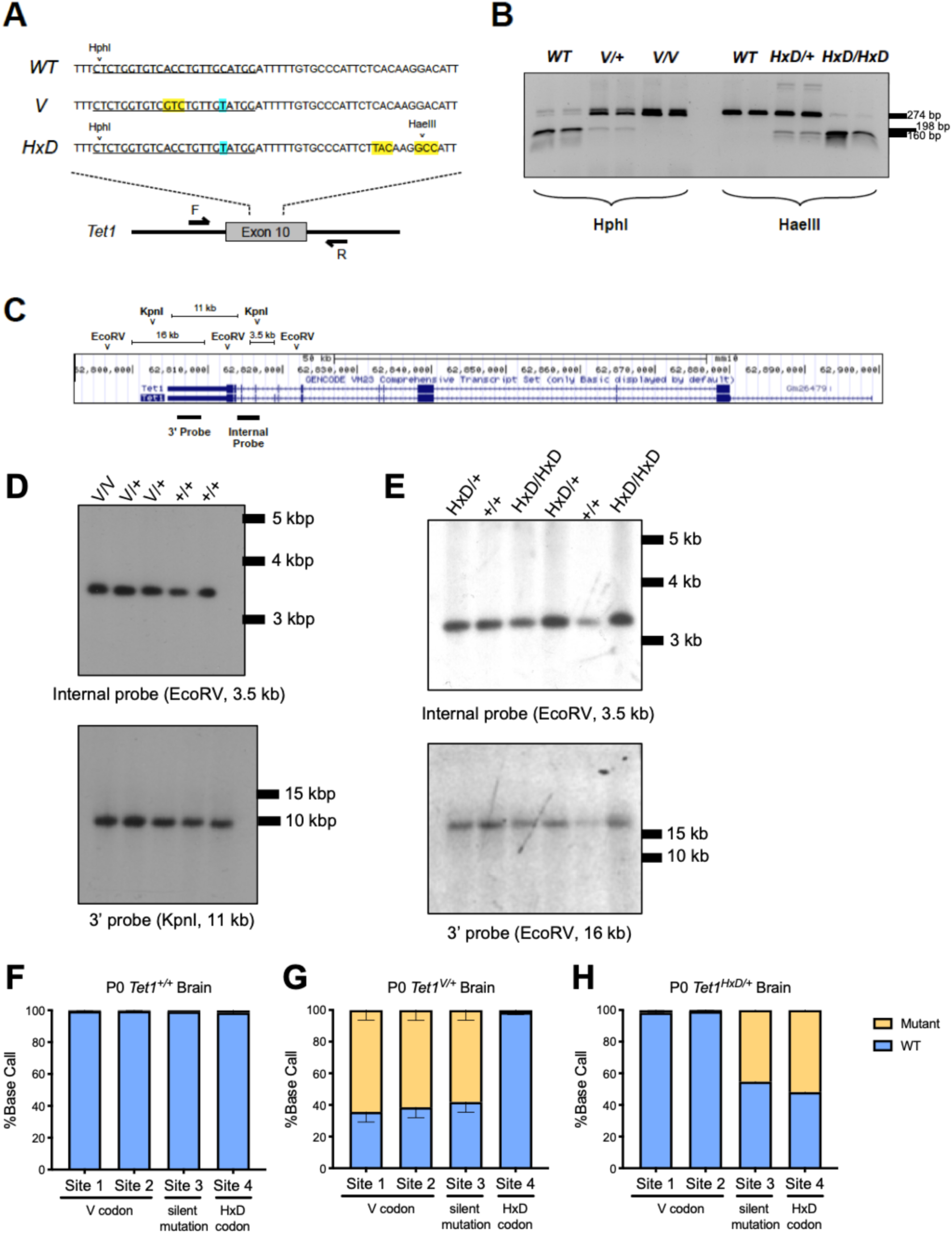
Additional validation of *Tet1-V* and *Tet1-HxD* mutant mouse lines. A) Exon 10 sequences for WT, *Tet1^V^*, and *Tet1^HxD^* alleles. Underlined sequence denotes the CRISPR gRNA target sites, while highlighted regions indicate non-synonymous (yellow) or silent (blue) mutations introduced by the HDR templates. Sequencing primers (F and R) amplify from flanking intronic regions to produce a 450 bp amplicon for RFLP analysis. B) Representative RFLP genotyping assay for *Tet1^V^* and *Tet1^HxD^*. C) Schematic depiction of restriction sites and radiolabeled probes used for *Tet1* Southern blots. D-E) Southern blot confirmation of *Tet1^V^* (D) and *Tet1^HxD^* E) mutant alleles with preserved fragment sizes. F-H) Allele specific RT-PCR followed by pyrosequencing for expression from *Tet1^V^* or *Tet1^HxD^* or WT allele in heterozygote PND0 brains.

**Supplemental Figure 2.**
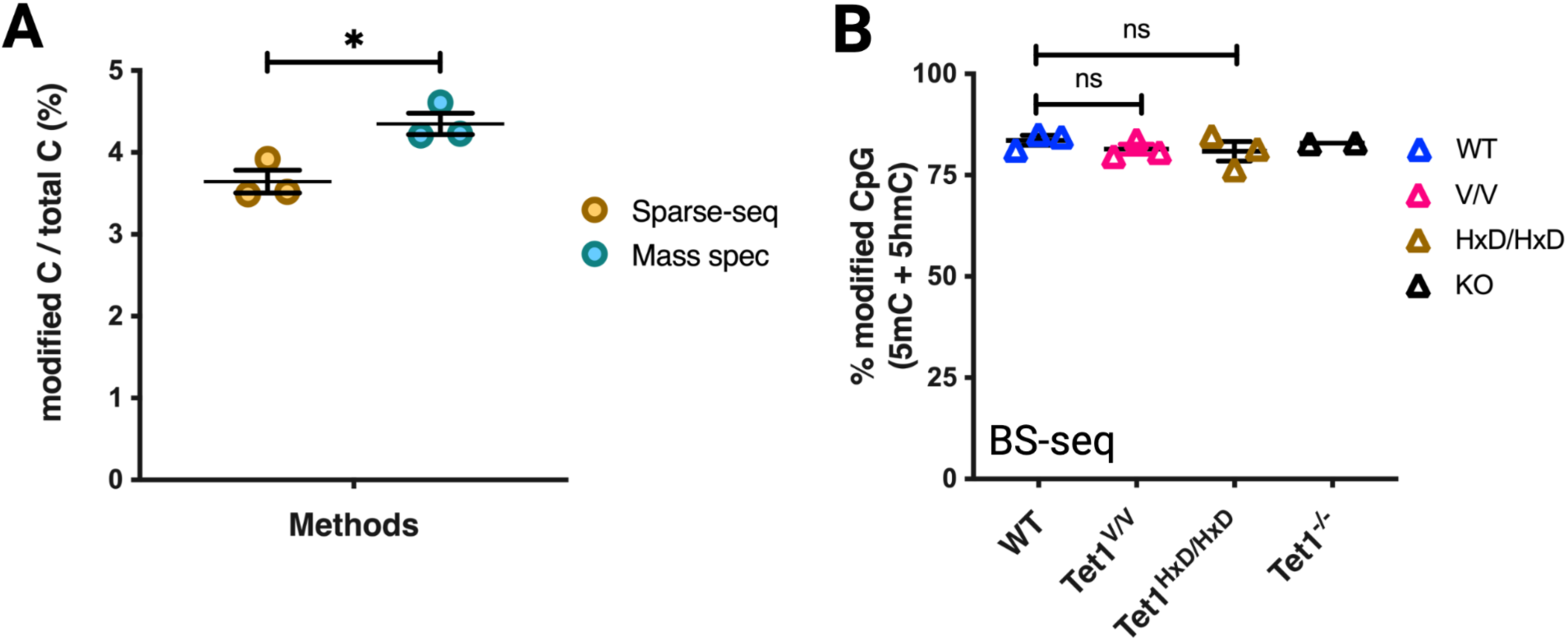
Validation of sparse-BS-seq to estimate modified cytosine levels. A) Sparse-BS-seq of 40-week-old cortex samples show comparable levels of modified C in all sequence contexts compared to mass spectrometry (mean ± SEM; n=3; t-test, *p-value < 0.05). B) Sparse-seq for total modified cytosine (5mC and 5hmC, BS-seq) in CpG context in adult testes show unchanged levels, globally, in mutants compared to WT.

**Supplemental Figure 3.**
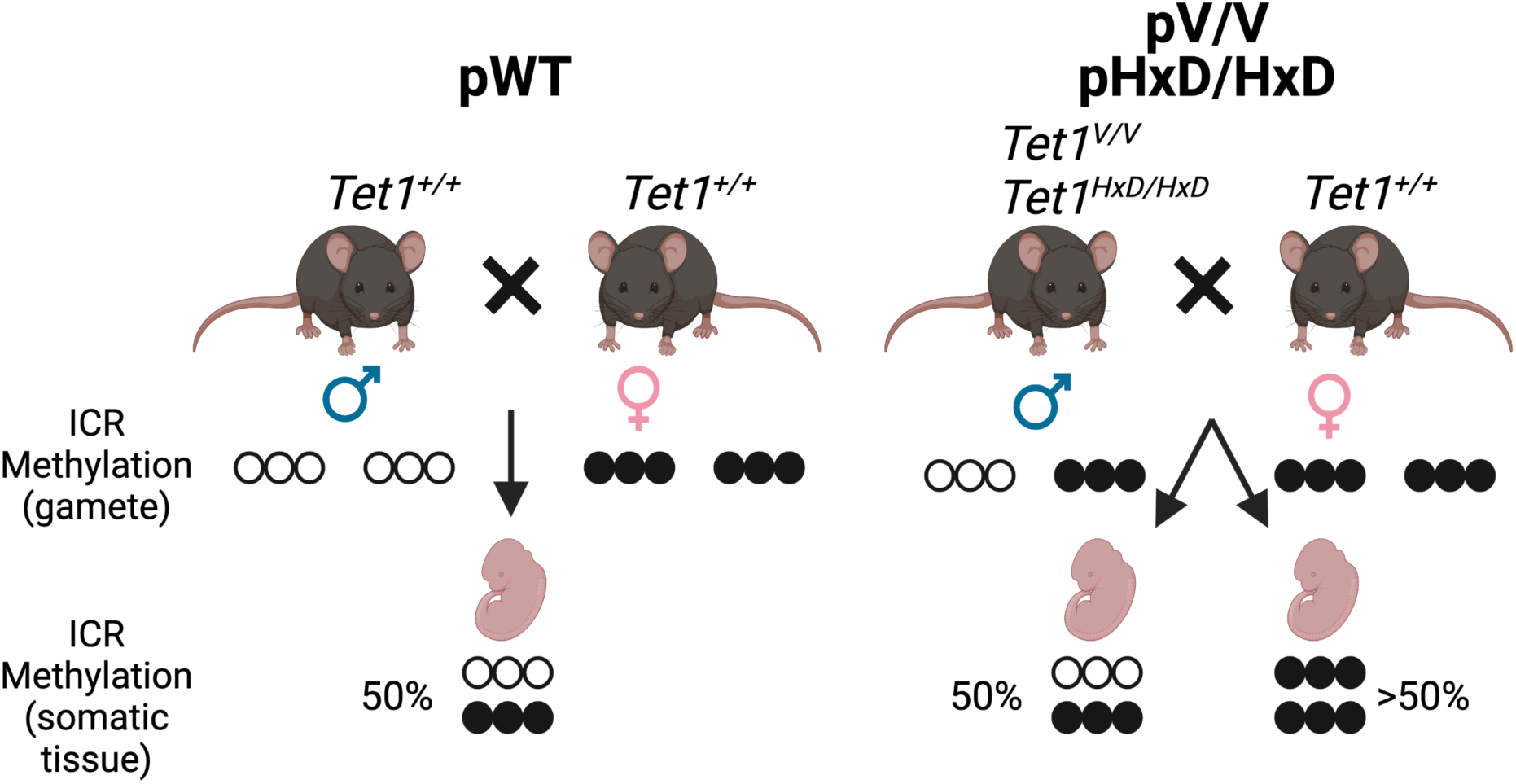
Breeding schemes used to test the heritability of imprinting defects in the germline of *Tet1-variant* mutant males. To generate pWT, WT male litter mates (*Tet1^+/+^*) are mated with C57BL/6J (B6) females. pVV or pHxD are generated by mating *Tet1^V/V^* or *Tet1^HxD/HxD^* males to B6 females, embryos are collected at gestational stage E10.5 or pups are collected at PND0. Hypothesized DNA methylation patterns for maternally methylated ICRs (that are normally unmethylated in the sperm) are indicated below each breeding scheme.

**Supplemental Figure 4.**
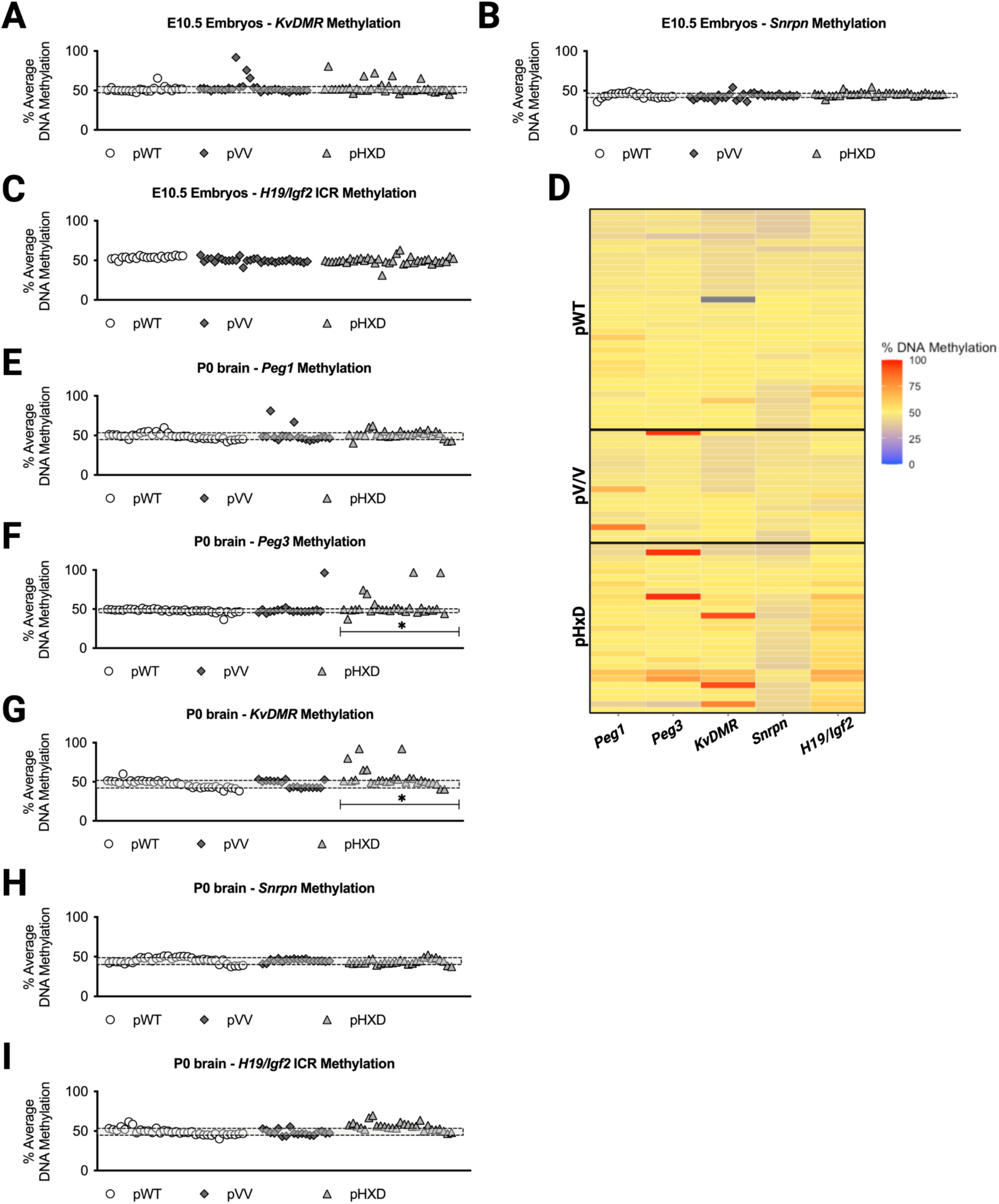
Additional characterization of ICR methylation defects in offspring of *Tet1 variant* mutant males. Percentage DNA methylation at maternally methylated *KvDMR* (A) and *Snrpn* (B) and a control paternally methylated ICR *H19/Igf2* (C). pWT n=22 embryos (3 litters), pVV n=31 (4 litters), pHxD n=37 (4 litters). D) DNA methylation levels at representative ICRs measured in the brain of individual PND0 offspring from *Tet1^+/+^*, *Tet1^V/V^* and *Tet1^HxD/HxD^* males are presented as a heatmap. Each row represents an individual PND0 of the indicated paternal genotype. Percentage DNA methylation at *Peg1* (E), *Peg3* (F), *KvDMR* (G), and *Snrpn* (H) and control *H19/Igf2* (H) ICRs as measured by pyrosequencing. pWT n=35 pups (5 litters), pVV n=18 pups (5 litters), pHxD n=28 pups (5 litters). *p<0.05, **p<0.01 Fisher’s exact test for frequency of hypermethylated embryo, shaded bars indicate average methylation of pWT embryos ± STDEV. Embryos with methylation above or below the shaded bar are considered hyper- or hypomethylated.

**Supplemental Figure 5.**
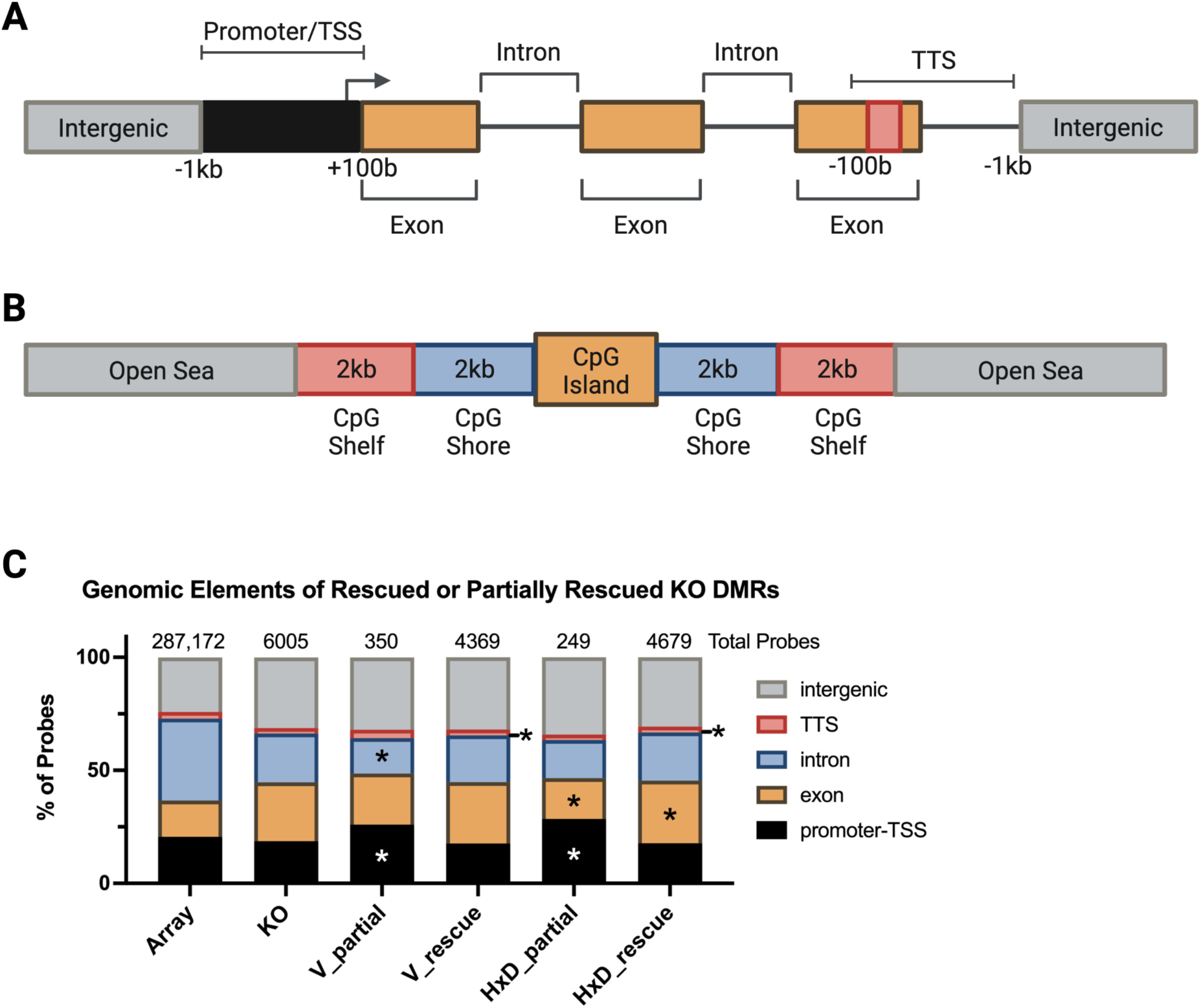
Genomic compartmental analysis for rescued DMRs in *Tet1*-catalytic mutants. A) Diagram depicting where promoter-TSS, exon, intron, TTS, and intergenic regions are defined in HOMER annotatePeaks basic annotation function. Promoter-TSS is defined from -1kb to +100bp of RefSeq annotated transcriptional start site and TTS is defined from -100bp to +1kb of RefSeq annotated transcriptional termination site. B) Diagram depicting CpG island annotation by R package AnnotationHub. CpG shores are defined as +/-2kb from the ends of the CpG island, excluding the CpG islands. CpG shelves are defined as +/-4kb from the ends of the CpG island, excluding the CpG islands and CpG shores. The remaining genomic regions are defined as open seas. C) Bar graph showing genomic localization of *Tet1^-/-^* DMRs that are either partially rescued or fully rescued in *Tet1^V/V^* or *Tet1^HxD/HxD^* sperm, as defined in Figure 3E-F. Bernoulli distribution test was conducted to compare genomic distribution of rescued or partially rescued probes to the genomic distribution of all differentially methylated probes (DMRs) in *Tet1^-/-^*. Partially rescued probes are more likely to be found in gene bodies (promoter-TSS, exon, and intron) while fully rescued probes are overrepresented at exon and TTS. *p-value < 0.05; two-sided Bernoulli distribution test as compared to genomic distribution *Tet1^-/-^* DMRs.

**Supplemental Figure 6.**
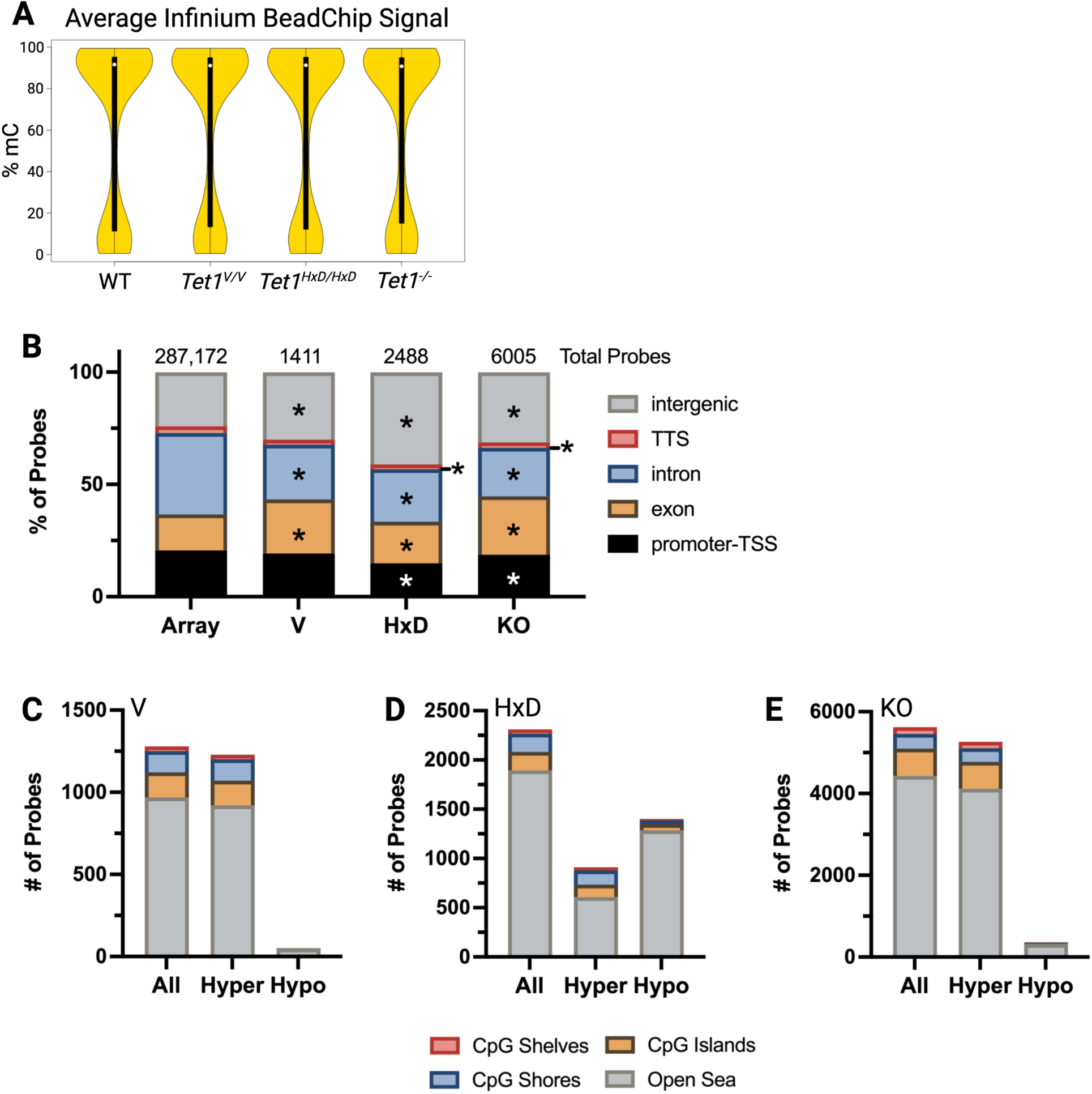
Genomic distributions of differentially methylated probes are distinct between *Tet1^-/-^*, *Tet1^V/V^*, and *Tet1^HxD/HxD^* sperm. A) Violin plot showing average methylation signal of all probes in *Tet1^-/-^*, *Tet1^V/V^, Tet1^HxD/HxD^* and WT sperm. B) Bar graphs showing genomic localization of DMRs from mutant sperm as annotated by HOMER. While intergenic and exonic regions are overrepresented as DMRs for the mutants, other regions are underrepresented relative to the array coverage. Total probes refer to total probes in the array or total probes that are significantly different in mutant sperm vs. WT. *p-value < 0.05; two-sided Bernoulli distribution test as compared to genomic distribution of all probes in the array. C-E) Bar graphs showing CpG density of DMRs in mutant sperm as annotated by R-package: annotatr.

**Supplemental Figure 7.**
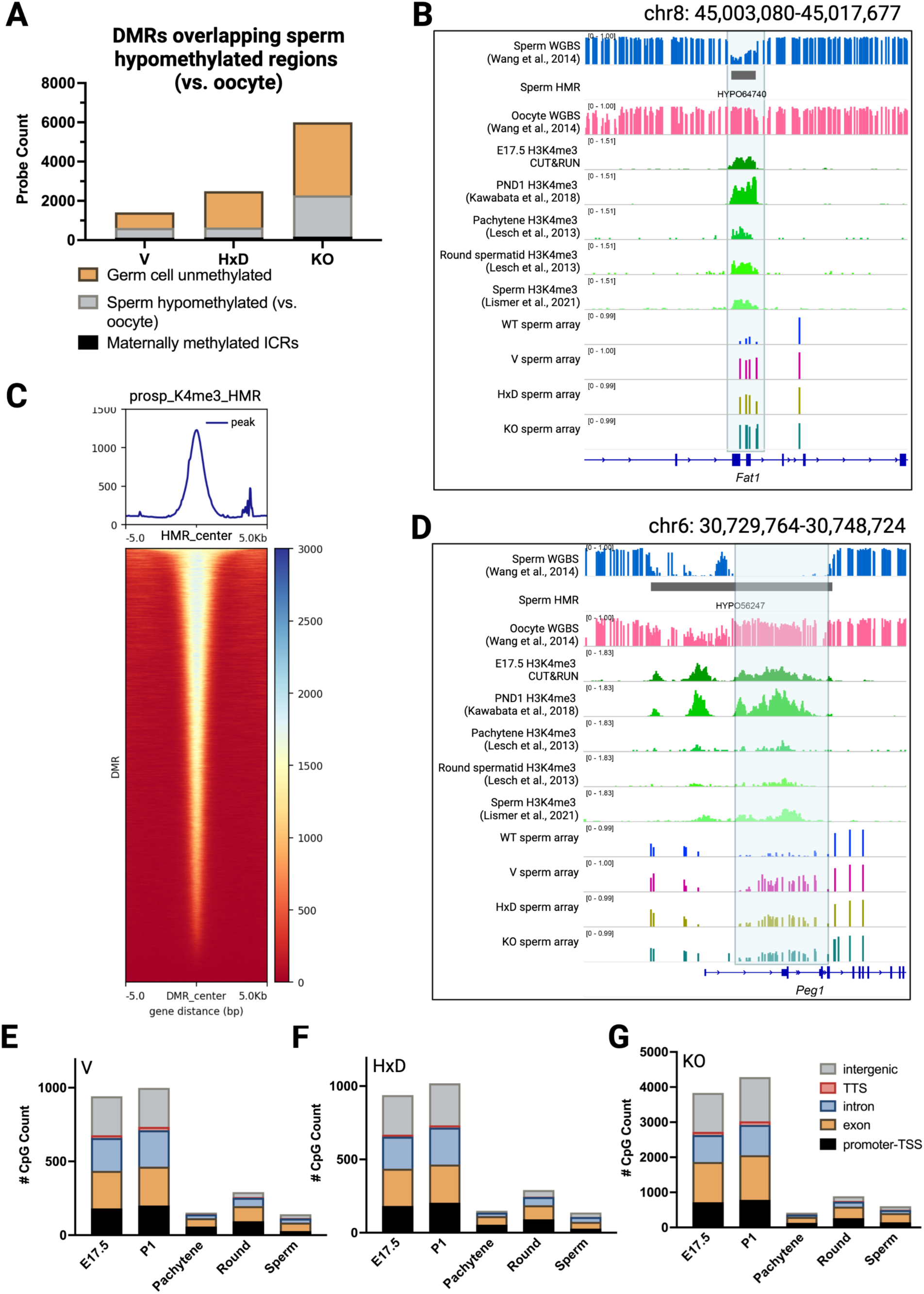
H3K4me3 enrichment at TET1-dependent sperm hypomethylated regions throughout spermatogenesis. A) Distribution of DMRs that overlap unmethylated regions in the sperm genome compared to the oocyte genome (GEO: GSE56697^72^). B, D) Genome browser view of an example locus (*Fat1)* and the canonical ICR *Peg1* (D) that are commonly hypermethylated in *Tet1^-/-^*, *Tet1^V/V^, Tet1^HxD/HxD^* sperm with overlap to the sperm hypomethylated region and H3K4me3 enrichment throughout spermatogenesis. C) Heatmaps and metapots of E17.5 prospermatogonia H3K4me3 enrichment centered on sperm HMRs, indicating majority of H3K4me3 signals at H3K4me3 were overlapping unmethylated regions in the sperm genome. Genomic distribution of differentially methylated probes in *Tet1^V/V^* (E), *Tet1^HxD/HxD^* (F), and *Tet1^-/-^* (G) sperm that overlap with H3K4me3 enrichment in E17.5 and PND1 prospermatogonia (DDBJ: DRA006633^35^), pachytene spermatocyte, round spermatid (SRA097278^73^, and sperm (GEO: GSE135678^74^).

## STAR Methods

### Key Resources Table

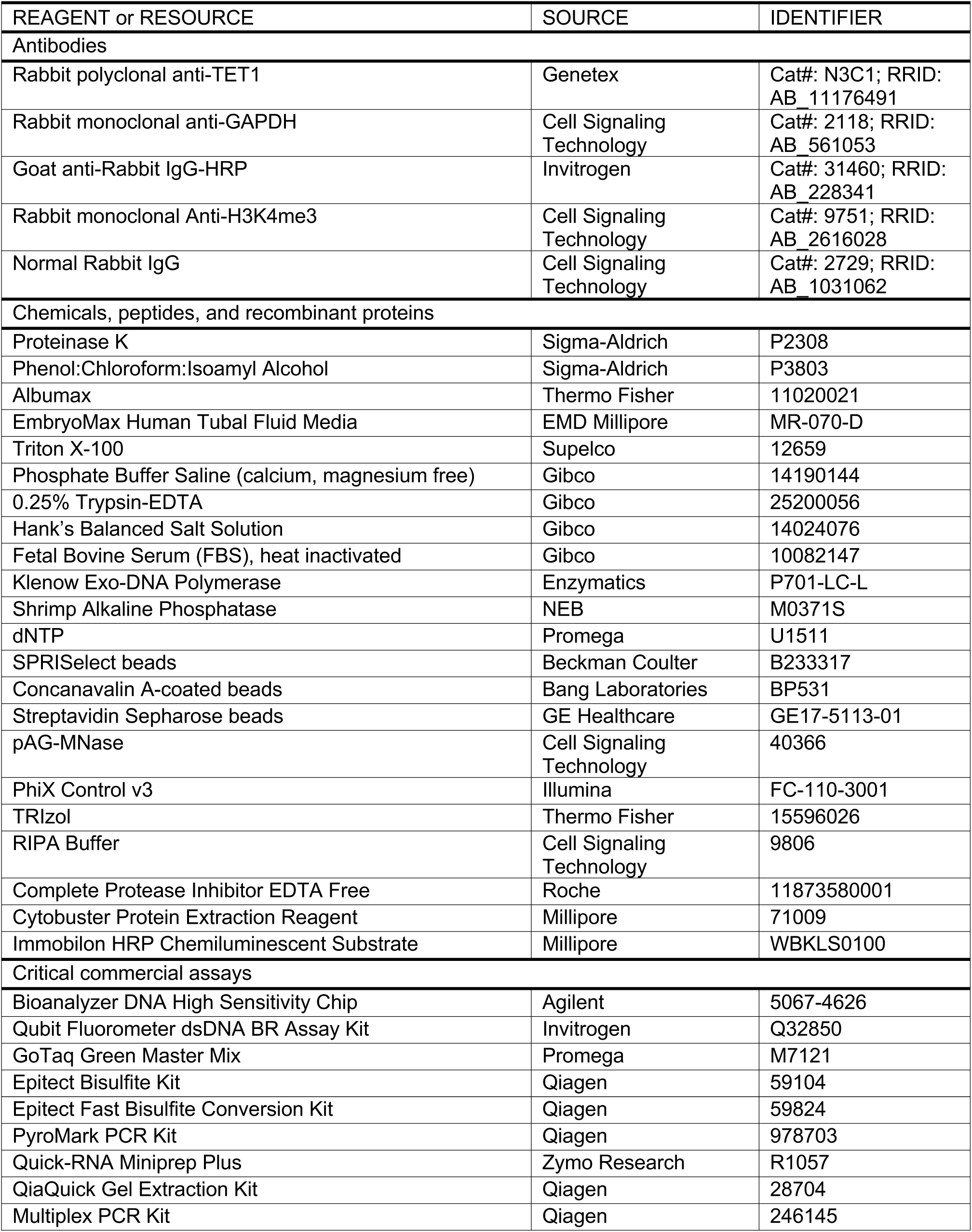
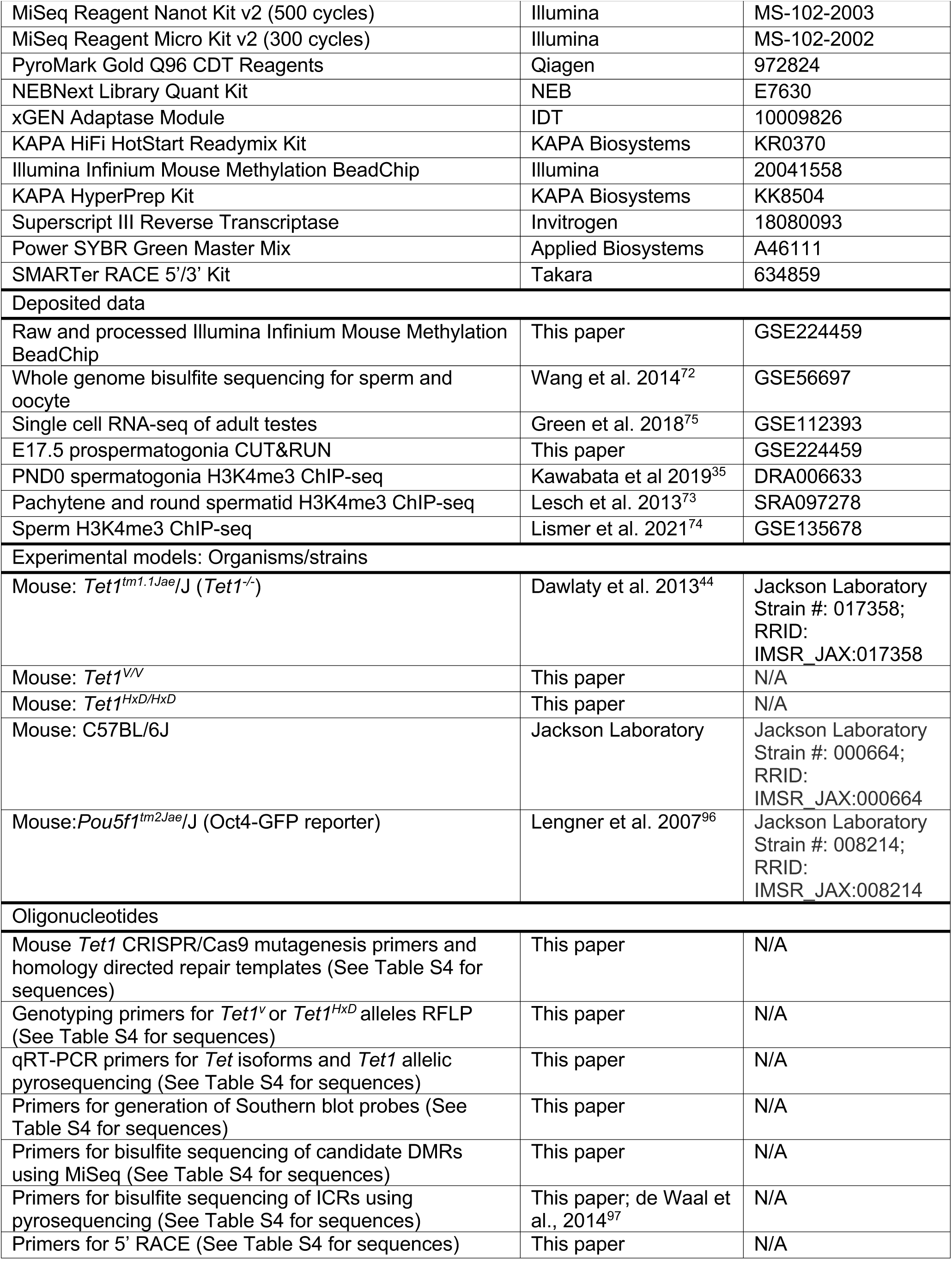
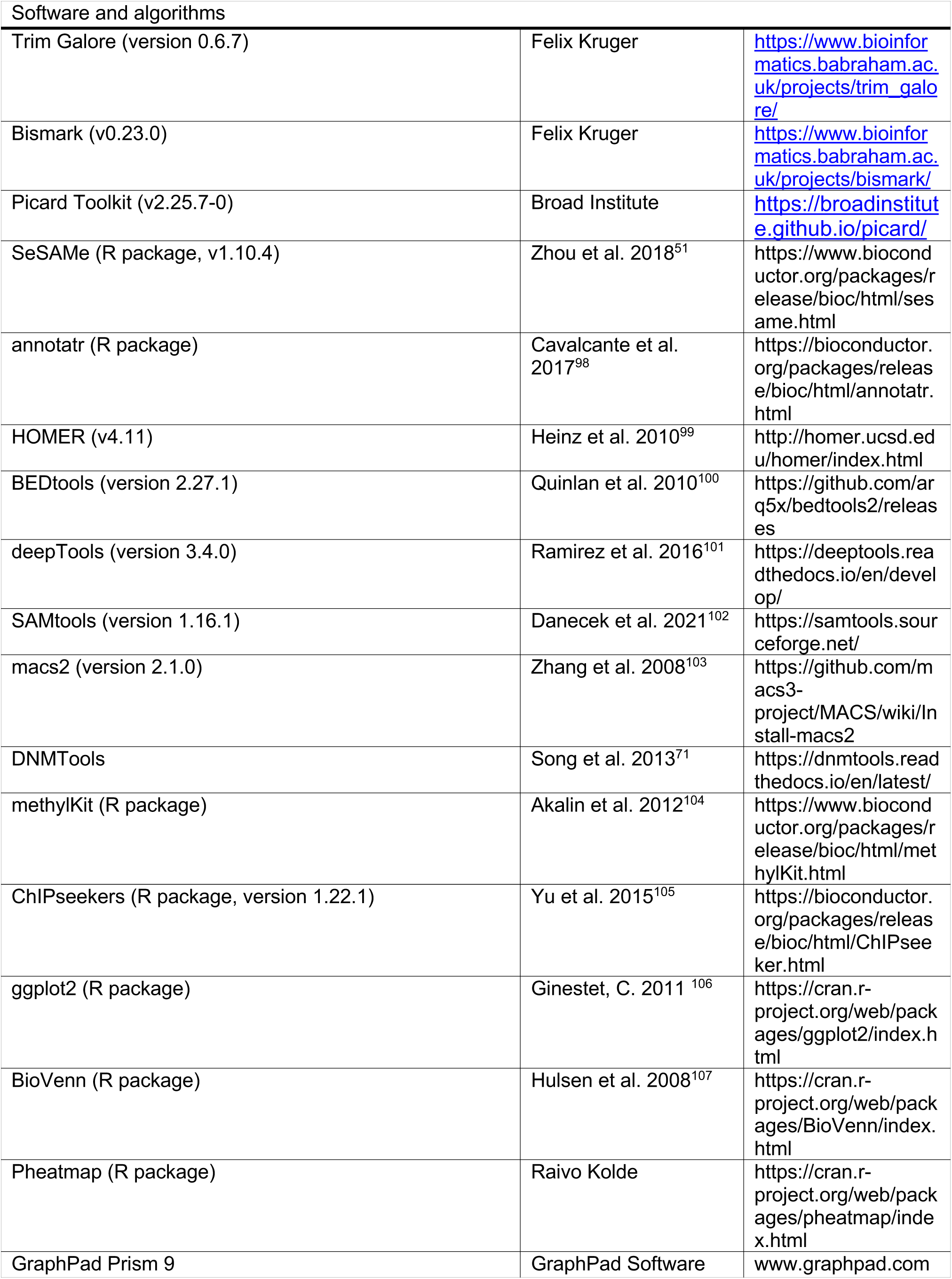

### Lead Contact

Further information and requests for resources and reagents should be directed to and will be fulfilled by co-Lead Contacts, Marisa S. Bartolomei and Rahul M. Kohli

### Materials Availability

All unique reagents and new mouse lines generated in this study are available from the Lead Contacts with a completed Materials Transfer Agreement.

### Data and Code Availability

The accession number for raw and processed Illumina Mouse Infinium Methylation BeadChip and CUT&RUN data generated in this paper is GEO: GSE224459. Accession numbers for existing, publicly available data are referenced as appropriate. Any additional information required to reanalyze the data reported in this paper is available from lead contacts upon request.

## EXPERIMENTAL MODEL AND SUBJECT DETAILS

### Mouse husbandry and maintenance

All experiments were approved by the Institutional Animal Care and Use Committee of the University of Pennsylvania (protocol number: 804211). Mice are housed in polysulfone cages within a pathogen-free facility with 12-12 h light-dark cycle and *ad libitum* access to water and standard chow (Laboratory Autoclavable Rodent Diet 5010, LabDiet, St. Louis, MO, USA). *Tet1* knockout^108^ (017358; B6; 129S4-*Tet1^tm1.1Jae^*/J) and *Oct4-GFP*^96^ (008214; B6; 129S4-*Pou5f1^tm2.Jae^*/J) were purchased from The Jackson Laboratory and were backcrossed for at least 10 generations to C57BL/6J (B6; The Jackson Laboratory, 000664) background. *Oct4-GFP* allele was maintained as homozygous in *Tet1* heterozygote breeders (*Tet1^+/-^*; *Oct4^GFP/GFP^*).

Mouse genomic DNA for genotyping using polymerase chain reaction (PCR) was isolated from ear punches as previously described^11^. Primers used for genotyping of *Tet1^-^*, *Tet1^V^*, *Tet1^HxD^*, *Oct4-GFP* alleles as well as sex genotyping are listed in Table S4. Timed mating was determined by visual detection of a vaginal sperm plug where E0.5 was taken to be 12.00h (noon) on the day the plug was observed. Visual staging of embryonic age was done at the time of dissection.

### Generation and validation of *Tet1^V^* and *Tet1^HxD^* mouse lines

Mutational insertions at exon 10 of endogenous *Tet1* allele to generate T1642V or H1654Y;D1656A amino acid substitutions within catalytic domain of TET1 were done using easi-CRISPR-Cas9 editing in the C57BL/6J (B6) and B6D2 (hybrid of B6 and DBA/2J) as previously described^109–111^. Briefly, single-strand homology-directed repair (HDR) donor templates carrying the appropriate nucleotide substitutions in *Tet1* exon 10 were synthesized as 4 nM Ultramers from Integrated DNA Technologies (see Table S4). The *Tet1* exon 10 guide RNAs (gRNAs) were amplified using synthesized oligos and the pX335 gRNA scaffolding vector, *in vitro* transcribed using MEGAshortscript T7 transcription kit (Ambion), and purified using MEGAclear Transcription Clean-up kit (Ambion). The purified gRNA (50 ng/uL), Cas9 mRNA (100 ng/uL), and HDR donor templates (100 ng/uL) were injected into the pronucleus of single-cell B6 x B6D2 embryos and transferred to pseudopregnant dams. The mosaic founders from the CRISPR injection were ∼75% B6 genetic background. Founders were screened by exon 10 restriction fragment length polymorphism (RFLP) using *HaeIII* enzyme to distinguish *Tet1^HxD^* allele and *HphI* enzyme to distinguish *Tet1^V^* allele from WT, of PCR amplified product flanking exon 10 (see Table S4). Genotypes of founders were further verified by Sanger sequencing. The targeted *Tet1* allele was validated in heterozygote and homozygote animals after backcrossing to B6 strain for at least 3 generations using Southern blot as previously described^112^, with restriction enzymes and probes indicated in Supplemental Figure 1 and Table S4. All mice included in this study had been backcrossed to B6 strain for at least 4 generations unless noted otherwise.

## METHOD DETAILS

### Tissue collection

#### Sperm

Adult male mice (>10 weeks of age) were housed with a sexually mature female for at least 5 days, and then isolated for at least 3 days. After euthanasia, the caudal epididymis was dissected and the epididymal sperm was collected on a needle and capacitated in EmbryoMax Human Tubal Fluid media (HTF, EMD Millipore) for 30 minutes at 37°C. Motile sperm were collected by removing the supernatant, spun down for 5 minutes at 650 xg and incubated for 15 minutes on ice with somatic cell lysis buffer (0.1% SDS, 0.5% Triton-X-100) to remove any non-sperm contaminants. Following treatment with somatic lysis buffer, the sperm were counted, spun down for 5 minutes at 10,000 xg and snap frozen for storage at -80°C until further processing.

### Embryonic germ cells

Embryonic *Oct4-GFP^+^* gonads were harvested from embryos at E12.5, E14. And E17.5. The gonads were dissected in calcium- and magnesium-free phosphate buffered saline (PBS, Gibco) and transferred into 500 μL of 0.25% Trypsin-EDTA (Gibco). Gonads were incubated in Trypsin-EDTA for 10 minutes at 37°C and quenched with equal volume of Hank’s Balanced Salt solution (HBSS, Gibco) containing 5% fetal bovine serum (FBS). To achieve single cell suspension, gonads were triturated using p1000 tips (Denville), followed by p200 tips (Denville) and 22G needle (BD Biosciences). The single cell suspension was centrifuge for 5 minutes at 650 xg and resuspended in 5% FBS in HBSS prior to sorting. GFP+ PGCs were sorted using FACSAria Fusion or FACS Jazz cell sorter (Becton Dickinson). For bisulfite mutagenesis, PGCs were snap frozen for storage at -80°C until further processing. For CUT&RUN, PGCs were immediately processed for permabilization and binding to concanavalin A beads.

### Somatic tissues

E10.5 whole-embryo was collected from timed-mating and immediately snap frozen for storage at -80°C until further processing. PND0 brain, tongue, and liver samples were collected following decapitation of neonates. Whole testis, liver, and cortex were dissected from male mice following euthanasia by CO2 asphyxiation. Tissues were snap frozen for storage at -80°C until further processing.

### Tissue homogenization and DNA extraction

Embryonic, neonatal, and adult tissues were digested in lysis buffer (50 mM Tris, pH8.0, 100 mM EDTA, 0.5% SDS) with proteinase K (180 U/mL; Sigma-Aldrich) overnight at 55°C. Sperm pellets were resuspended in sperm lysis buffer (20 mM Tris-HCl, pH8.0, 200 mM NaCl, 20 mM EDTA, 4% SDS) with the addition of 5 μL of β-mercaptoethanol and proteinase K (180 U/mL) at 55°C overnight. Genomic DNA was isolated using Phenol:Chroloform:Isoamyl Alcohol (25:24:1; Sigma-Aldrich) and ethanol precipitation and resuspended in TE buffer (10 mM Tris-HCl pH8.0, 0.5 mM EDTA).

### Bisulfite mutagenesis

2 μg of genomic DNA was bisulfite treated using the EpiTect Bisulfite Kit (Qiagen) and eluted in 20 μL of 1:10 of the supplied EB buffer. Snap frozen PGC pellet was directly lysed using the LyseAll Lysis Kit (part of the EpiTect FAST Bisulfite Conversion Kit, Qiagen) and was bisulfite treated using the standard Epitect Bisulfite reagent mix following the low-input protocolPGC bisulfite-treated DNA was resuspended in 20 μL of 1:10 of the supplied EB buffer.

### Library preparation for bisulfite sparse sequencing

Whole genome BS libraries preparation was adapted from Luo et al.^113^ using xGEN Adaptase module (Integrated DNA Technology) following the workflow for single-cell Methyl-Seq (snmC-Seq), which includes random priming step, per manufacturer’s protocol. Additional components for random priming step are as followed: Klenow Exo-DNA Polymerase at 50 U/μL supplied with Blue Buffer (Enzymatics), Exonuclease I at 20 U/μL (Enzymatics), Shrimp Alkaline Phosphatase (NEB), 10 mM dNTP (Promega). Following random priming step, samples were eluted in 10 μL Low EDTA TE (included in the xGEN Adaptase module) and proceeded with Adaptase reaction per manufacturer’s protocol. To determine cycle numbers for enrichment PCR, 1 μL of library from Adaptase reaction was used to run qRT-PCR with the following condition, 0.5 uL of 10uM custom Illumina I7 and I5 primers to accommodate stubby adapter tails on the random primers (final concentration of 0.5 μM and all unique dual index primer sequences are listed in Luo et al.^113^), 7 μL of 2x NEBNext Library Quant Master Mix with 1:100 low ROX (NEB), and 1.5 μL ddH2O. Indexing PCRs were done with 3 cycles less than the determined qRT-PCR cycle threshold (Ct) using KAPA HiFi HotStart ReadyMix (KAPA Biosystems) with final custom Illumina I7 and I5 concentrations at 1 μM. Amplified libraries were cleaned using two rounds of 0.8X SPRISelect beads and eluted in 13 μL of EB Buffer. Libraries were quantified using NEBNext Library Quant Kit and libraries sizes were determined using Bioanalyzer High Sensitivity DNA Kit (Agilent). Indexed libraries were pooled and sequenced on an Illumina MiSeq using a MiSeq Reagent Kit v2 (150×150; Illumina) with 10% PhiX spike-in to achieve ∼65,000 aligned reads per library.

### Locus specific DNA methylation analysis using pyrosequencing or targeted next-generation bisulfite-sequencing

Pyrosequencing PCRs and sequencing reactions for ICR methylation (*H19/Igf2, Peg1, Peg3, KvDMR*, and *Snrpn)* were described by SanMiguel et al^11^. Primers are listed in Table S4. Targeted DNA methylation analyses using next-generation sequencing were modified from IMPLICON protocol^114^. For assay design, genomic DNA sequences of the regions of interest were obtained from UCSC Genome Browser and imported into MethPrimer^115^ or BiSearch^116^ to identify primer pairs with optimal amplicon size of 300 bp with a minimum of 5 CpGs. Primers are listed in Table S4. Stubby Illumina adapter sequences were added to the forward (ACACTCTTTCCCTACACGACGCTCTTCCGATCT) and reverse (GTGACTGGAGTTCAGACGTGTGCTCTTCCGATCT) primers. 1 μL of bisulfite treated DNA was amplified for the regions of interest using PyroMark PCR kit (Qiagen) with final primer concentration of 0.4 μM. Amplicons of similar sizes (+/-50 bp) were pooled for column purification (Thomas Scientific) and eluted in TE buffer. 25 ng of pooled amplicons were loaded into indexing PCR using Multiplex PCR kit (Qiagen). Indexed reactions were purified using SPRIselect beads (0.9x; Beckman Coulter), eluted in TE buffer and quantified using Qubit Flourometer dsDNA BR Assay Kit (Invitrogen). Indexed libraries were pooled and sequenced on an Illumina MiSeq using a MiSeq Reagent Nano Kit v2 (250×250; Illumina) with 10% PhiX spike-in.

### Western blot

Testes were homogenized by mortar and pestle in 1x RIPA buffer (Cell Signaling Technology) supplemented with Complete Protease Inhibitor (Roche). Tissue lysates were incubated on ice for 20 minutes, followed by centrifugation at 13,000 xg for 5 minutes at 4°C. Supernatant was collected and the protein concentration was quantified using bicinchoninic acid protein assay (BCA assay; Pierce, Thermo Scientific). 50 μg of protein lysate was denatured and were run on a 4-12% SDS-PAGE gel. The gel was transferred onto a PVDF membrane at 250 mA for 120 minutes, and then blocked for 1 hour at RT with shaking in 5% non-fat dry milk in TBS-Tween (TBS-T). For full-length TET1 detection, membranes were probed with 1:1000 rabbit anti TET1 (N3C1, Genetex) overnight at 4°C. For loading control, membranes were probed with 1:10,000 rabbit anti-GAPDH (2118, Cell Signaling Technology) overnight at 4°C. Membranes were washed 3x in TBS-T following primary antibody incubation, and probed with 1:20,000 goat anti-rabbit IgG HRP secondary antibody (Invitrogen) for 1 hour at RT. Membranes were developed using Immobilon Western Chemiluminescent HRP Substrate and imaged on an Amersham Imager 600.

### Genome-wide DNA methylation profiling using Infinium Mouse Methylation BeadChip

1000 ng of bisulfite-treated sperm DNA was loaded onto Illumina Infinium Mouse Methylation-12v1-0 BeadChip (llumina) and was ran on an Illumina iScan System (Illumina) per manufacturer’s standard protocol. The samples were processed at the Center for Applied Genomics Genotyping Core at the Children’s Hospital of Philadelphia. Biological replicates for each genotype are as followed, *Tet1^+/+^* n = 8, *Tet1^V/V/^* n = 10, *Tet1^HxD/HxD^* n= 8, *Tet1^-/-^* n = 10.

### E17.5 Prospermatogonia CUT&RUN

E17.5 *Oct4-GFP^+^* prospermatogonia were collected using FACS as detailed above. Cells were hold on ice until CUT&RUN processing. Gonads from several embryos were pooled and CUT&RUN was done on 130,000 freshly sorted cells per motif (H3K4me3 and IgG) as previously described^117^. Sorted cells were spun down at 600 xg for 3 minutes at room temperature, and washed three times with 1.5 mL of Wash buffer (20 mM HEPES pH 7.5, 150 mM NaCl, 0.5 mM spermidine, with 1x Roche complete protease inhibitor EDTA Free). Cells were bound to concanavalin A-coated magnetic beads (Bangs Laboratories) pre-washed in binding buffer (20 mM HEPES (pH 7.5), 10 mM KCl, 1 mM CaCl2, 1 mM MnCl2) for 10 minutes at room temperature. Cells were resuspended with H3K4me3 (Cell Signaling Technology 9751) or rabbit IgG antibodies (Cell Signaling Technology 2729) at a final dilution of 1:100 in 200 μL of antibody binding buffer (same as Wash buffer with 0.05% digitonin and 2 mM EDTA) and incubated overnight at 4°C. Cells were washed twice in Wash buffer with 0.05% digitonin, and incubated with pAG-MNase fusion protein (final concentration 700 ng/mL) for 1hr at 4°C. Following two washes in Wash buffer with 0.05% digitonin, samples were resuspended in Incubation buffer (3.5 mM HEPES (pH 7.5), 10 mM CaCl2, 0.05% digitonin) to activate cleavage on ice for 5 minutes. 2xSTOP solution (170 mM NaCl, 20 mM EGTA, 0.05% digitonin, 50 μg/mL RNase A, 25 μg/mL Glycogen, 2 pg/mL yeast chromatin spike-in) was added to quench the reaction. To release antibody bound fragments, samples were incubated at 37°C for 30 minutes and supernatant was isolated from the magnet-bound beads. 2 μL of 10% SDS and 2.5 μL of 20 mg/mL Proteinase K were added to the suprnatang and samples were incubated at 50°C for one hour to digest any protein, and DNA was isolated using Phenol:Chloroform extraction twice. DNA pellets were resuspended in 30 μL of Tris EDTA buffer (1mM Tris HCl, pH 8; 0.1 mM EDTA). CUT&RUN library was prepared using KAPA HyperPrep Kit (KAPA Biosystems) following the manufacturer’s protocol. Libraries were amplified for 18 cycles using KAPA HiFi Hot Start Ready Mix (Roche). Amplified libraries were cleaned using 1x KAPA Pure Beads Libraries were quantified using NEBNext Library Quant Kit and libraries sizes were determined using Bioanalyzer High Sensitivity DNA Kit (Agilent).

### RNA extraction, reverse transcription, qRT-PCR, and pyrosequencing for allelic expression analysis

For adult testes or PND0 livers, tissue lysates were divided in half by volume and added to TRIzol reagent (Thermo Fisher Scientific). Chloroform was added to achieve phase separation, followed by recovery of the aqueous phase. Equal volume of ethanol was added to the aqueous phase and RNA was bind to Zymo Research RNA miniprep column. RNA purification was done following Quick-RNA Miniprep Plus Kit as specified by manufacturer’s protocol including in column DNAseI treatment (Zymo Research). RNA quantity was determined by NanoDrop ND-1000 spectrophotometer (Thermo Fisher Scientific). RNA samples were reverse transcribed with Superscript III Reverse Transcriptase (Invitrogen) according to the manufacturer’s protocol. Quantitative real-time PCR (qRT-PCR) was performed using Power SYBR Green Master Mix (Applied Biosystems) on a QuantStudio 7 Flex Real-Time PCR system. Relative expression levels were determined using the Pffafl method normalized to the housekeeping gene *Nono.* To assess expression of *Tet1^HxD^* or *Tet1^V^* allele in heterozygote or homozygote PND0 brain, modified protocol for pyrosequencing for imprinting expression (PIE) was used as previously described in^111^ . 10 ng of cDNA was amplified using Pyromark PCR kit (Qiagen) with final primer concentration of 0.4 uM, in which the reverse primer was biotinylated. 4 uL of PCR product was sequenced on the Pyromark Q96 MD Pyrosequencer (Biotage, AB), using PyroMark Gold Q96 CDT Reagents (Qiagen), and Streptavidin Sepharose beads (GE Healthcare). Quantification of allele-specific expression was performed using Pyromark Q96 MD software based on the presence of a SNP (introduced in CRISPR mutagenesis) in the cDNA amplicon. Primers are listed in Table S4.

### 5’ rapid amplification of cDNA ends (RACE)

SMARTer RACE 5’/3’ Kit from Takara Bio was used following the manufacturer’s instructions. 3’ gene specific primers (Supplemental Table 4) were designed downstream of the hypothesized DMRs using Primer BLAST with parameters specified by SMARTer RACE kit (23-28 nucleotides long, 50-70% GC content, with Tm > 70°C) with GATTACGCCAAGCTT-overhang on the 5’ ends to allow for cloning into the provided pRACE vector. Reverse transcription was done on 1 μg of total RNA from adult testes. PCR condition modifications to amplify RACE products are as followed using SeqAmp DNA Polymerase (Takara Bio): *Fat1* (TA: 68°C and extension of 6 minutes) and *Dyrk2* (TA: 65°C and extension of 3 minutes). RACE products were gel isolated using Qiaquick Gel Extraction Kit (QIAGEN) and cloned into pRACE vector. Sanger sequenced products were subjected to BLAT alignment to mm10 genome.

## QUANTIFICATION AND STATISTICAL ANALYSIS

Statistics were performed using GraphPad Prism or R. Comparison of > 2 independent groups were performed using one-way ANOVA followed by Tukey’s post-hoc test. Fisher’s exact test was performed to determine significance of the frequency of hypermethylated F1 offspring of *Tet1* catalytic mutant males. Bernoulli distribution test was conducted to determine distribution of DMRs genomic compartment as compared to genomic distribution of all probes in the array. Statistical significances are denoted by different letters or asterisks in the graph. Information on statistical tests performed, exact values of n, and degrees of significance are provided in the figure legends.

### Analyses of sparse-seq and targeted next-generation bisulfite sequencing

Sequenced reads were trimmed using Trim Galore (version 0.6.7 https://www.bioinformatics.babraham.ac.uk/projects/trim_galore/) in paired-end mode. For sparse-seq, in addition to low quality bases and adaptors, 15 bps were removed from 5’ end of read 1 and 30 bps were removed from 5’ end of read 2 to remove synthetic sequences introduced by Adaptase Module random priming step. Trimmed sequenced reads were aligned to mouse mm10 genome with Bismark (v0. 23.0 https://www.bioinformatics.babraham.ac.uk/projects/bismark/) in paired-end mode. The following Bismark parameters were used to align sparse-seq reads: `--score_min L,0,-0.6 –non_directional`. Reads were deduplicated using Picard Toolkit (v2.25.7-0 https://broadinstitute.github.io/picard/) `MarkDuplicates.` Non-deaminated reads were filtered out based on the presence of ≥ 3 consecutive instances of non-CG methylation (Bismark function: `filter_non_conversion`; parameters`: --paired --threshold 3 –-consecutive`). Bedgraph files were prepared using Bismark Methylation Extractor function to calculate percent methylation at each CpG and global modified Cytosine levels (in CpG, CHG, CHH, and unknown context). For locus specific analysis, percent methylation at each CpG was calculated from at least 30x coverage.

### Analyses genome-wide DNA methylation profiling

Processing of raw IDAT files was done as previously described by Vrooman et al.^118^ using SeSAMe R Package^51^ (v1.10.4) and the MM285 array manifest file (vM25) to obtain methylation β-values (`getBetas` function). Probes that did not pass SeSAMe’s quality control pOOBAH approach for signal-to-background thresholding are masked with NA^51^. To determined differentially methylated probes, we included only CG probes with no NA values in all biological replicates, totaling in 218,483 probes out of 287,172 that are printed on the array. We first used SeSAMe `DML` (Differential Methylation Locus) function which models the DNA methylation levels using mixed linear model, a supervised learning framework that identifies CpG loci whose differential methylation is associated with known co-variates (i.e. sample genotype in our experiment)^51^. Only CG probes, as defined in “Infinium Mouse Methylation v1.0 A1 GS Manifest File.bpm”^47^, were included in F-test that is conducted as part of SeSAMe `DML` function and multiple-testing adjustment. To finalize the lists differentially methylated probes for each genotype compared to *Tet1^+/+^* was done using SeSAMe `DMR` function with false discovery rate cut off of 5% (Seg_Pval_adj < 0.05) and minimum difference of 10% between the mean of WTs and mutants. We used SeSAMe built in knowYourCG tool to test enrichments (`testEnrichment`) for probe design groups, chromHMM chromatin states, and transcription factor binding sites on differentially methylated probes for each mutant genotype compared to WT.

All downstream analysis was conducted using the mm10/GRCm38 mouse genome assembly. Differentially methylated regions (DMRs, each DMR corresponds to one array probe) were assigned to nearby genes and prioritized to genomic features based on proximity using HOMER `annotatePeaks.pl` function^99^. We calculated Bernoulli distribution of DMRs’ genomic features distributions as compared to the genomic features distribution of all probes in the array. CpG densities of DMRs were assigned using the annotatr R package^98^. Venn diagrams were generated using the R package BioVenn^107^. Partially supervised clustering was conducted on DMRs of all genotype to determine hyper- or hypomethylated signatures using the R package pheatmap with `average` clustering method. To assess rescue or partial rescue by TET1^V^ or TET1^HxD^, DMRs that were found in *Tet1^-/-^* were assessed in *Tet1^V/V^* or *Tet1^HxD/HxD^* samples. DMRs were classified as partially rescued if the FDRs were less than 0.05 in *Tet1^V/V^* or *Tet1^HxD/HxD^* but mean differential methylation levels between *Tet1^V/V^* or *Tet1^HxD/HxD^* samples and *Tet1^+/+^* were less than 10%. DMRs were classified as rescued if the FDRs were greater than 0.05 in *Tet1^V/V^* or *Tet1^HxD/HxD^* samples compared to *Tet1^+/+^*. Volcano plots were made using the R package ggplot2^106^. DMRs were examined for overlap with sperm HMRs, H3K4me3 ChIP-seq or CUT&RUN peaks for each stages of spermatogenesis using BEDtools^100^ (version 2.27.1) intersect. To generate heatmaps for H3K4me3 signals at DMRs, DMRs were first binned into 250 bp non-overlapping windows using BEDtools merge function, heatmaps and metaplots were generated using deepTools^101^(version 3.4.0).

### Analyses of ChIP-seq and CUT&RUN

Publically deposited ChIP-seq (DRA006633^35^, SRA097278^73^, GSE135678^74^) and CUT&RUN fastq files were trimmed using Trim Galore (version 0.6.7, default parameters). ChIP-seq and CUT&RUN reads were aligned to mouse mm10 reference genome using Bowtie2 (version 2.5.0, parameters: `--N 1 –sensitive -local`). Following alignment, low quality reads (QMAP ≤ 10) and non-primary alignments were removed using SAMtools^102^ (version 1.16.1, `view -q 10 -F 256`). Duplicated reads were removed using SAMtools `rmdup` with default parameter and mitochondrial reads were removed using `grep` function. Alignment BAM files were converted to BED files and blacklisted regions^119^ were remove using BEDtools (version 2.27.1). bigwig files normalized to count per million (CPM) were prepared using deepTools (version 3.5.1, function: `bamCoverage`, parameters: `-bs 1 --normalizeUsing CPM`). Peak calling was performed using Model-based Analysis of ChIP-Seq^103^(macs2, version 2.1.0; parameters: `--qvalue 0.01`).

### Identification of sperm hypomethylated regions

Sperm hypomethylated regions (HMRs) were determined using `hmr` function of DNMTools^71^, which uses hidden Markov model approach to identify methylation canyon in a supplied whole genome bisulfite sequencing methylation call data set (GEO: GSE56697^72^). In total, 76227 HMRs were identified in sperm WGBS data set. To identify hypomethylated regions in the sperm genome as compared to the oocyte genome, we used the R package methylKit^104^ `calculateDiffMeth` function.

### Processing of testis single cell RNAseq (scRNAseq)

Normalized gene expression matrices for adult mouse scRNA-seq spermatogenic germ cell clusters are publicly available (GSE112393^75^). DMRs were matched to their nearest annotated gene using the annotatePeaks function of the Rpackage ChIPSeeker^105^ (version 1.22.1; parameter: `TxDb=TxDb.Mmusculus.UCSC.mm10.knownGene`). Boxplots depicting normalized expression for DMR-linked genes vs. background gene expression for germ cell clusters were prepared using ggplot2^106^.

### BioRender

Figure 5B, Figure 7, Supplemental Figure 3, and Supplemental Figure 5A-B are created using BioRender.com

